# Social ties are maintained through sensory adaptations in green chromides

**DOI:** 10.64898/2026.11.03.763611

**Authors:** Chena Desai, Amit Babu, Rashmita Ganguly, Krishna B.S. Swamy, Ratna Ghosal

## Abstract

Sensory systems exhibit considerable plasticity in response to environmental changes, yet the functional implications of such plasticity remain poorly understood, particularly regarding how changes in visual gene expression influence social behaviour. To address this gap, we investigated how visual sensory plasticity relates to conspecific preferences in green chromides (*Etroplus suratensis*), an Indian cichlid that relies on vision for social preferences towards conspecifics. We first characterized retinal opsin and chromophore-related genes through transcriptomic analysis and subsequently examined behavioural and molecular responses under sequential white- and blue-light environments. We quantified visually mediated conspecific preferences, and the expression of rod and cone opsins and the chromophore-related gene Cyp27c1 at regular intervals. Green chromides retained stable conspecific preferences despite contrasting light environments. Opsins exhibited plasticity, with long wavelength-sensitive opsin (LWS) increasing under blue light, whereas short wavelength-sensitive opsins remained unchanged. Additionally, Cyp27c1 expression declined under blue light, suggesting a potential shift towards increased A1 chromophore utilization. Together, these findings suggest that green chromides undergo multiple molecular modifications under contrasting light environments, whereas maintaining stable social behaviours, demonstrating the cross-talk between the two is not straightforward. Overall, these results highlight that unraveling the underlying proximate mechanisms of behavior is highly intricate, involving multiple pathways, starting from perception (at the retinal level) to post-processing of the stimulus.

## 1 Introduction

Animal sociality is contingent upon the accurate perception of, and response to, environmental and social stimuli (Ward and Webster, 2016; Roberts and Roberts, 2020). When environmental conditions fluctuate, organisms often make adaptive adjustments in behavioral strategies, sensory processing, or both, to maintain ecological fitness (Bernhardt et al., 2020; Candolin and Wong, 2012; Canale and Henry, 2010). Although a substantial body of empirical literature confirms that animals exhibit significant plasticity in both behavioral and sensory domains to navigate heterogeneous environments, these faculties are traditionally investigated in isolation (Candolin and Wong, 2012; Canale and Henry, 2010). Crucially, social interactions are constrained by an animal’s *umwelt*, the species-specific perceptual reality dictated by its unique sensory limitations, which modulates the reception of ecological and social cues (Prior et al., 2021). Given that sensory mechanisms and social behavior are functionally coupled in mediating environmental interaction, a comprehensive understanding of adaptive strategies requires a holistic analysis of this socio-sensory network across varying conditions.

This paradigm is essential to visual ecology, as visual systems are fundamental to critical life-history behaviors, including foraging, mate selection, predator avoidance, navigation, and group formation (Lazavera et al., 2010; Ronald et al., 20212; Moore et al., 2015). While substantial evidence exists for high levels of visual sensory plasticity in response to environmental fluctuations, the functional consequences of this plasticity, specifically its influence on social behaviors, remain insufficiently explored (Nys et al., 2015, Bertinetti and Torres-Dowdall, 2025). In this context, teleosts (ray-finned fishes) represent an exceptionally suitable model, as they maintain considerable sensory and neural plasticity throughout adulthood, including ongoing neurogenesis in key visual centers such as the optic tectum (Bertinetti and Torres-Dowdall, 2025; Ebbesson and Braithewaite, 2012). Furthermore, the inherent variability of aquatic light environments, shaped by fluctuations in depth, turbidity, and organic matter, provides a robust framework for investigating the mechanisms of visual adaptation and their subsequent impact on behavioral performance (Carleton and Kocher, 2001).

In teleosts, visual function is governed by retinal opsin proteins located within rod and cone photoreceptors, which facilitate scotopic (low-light) and photopic (bright-light) vision, respectively (Carleton et al., 2020). These fishes possess a diverse suite of opsin classes, including RH1 for dim light, alongside SWS1, SWS2, RH2, and LWS for color discrimination, which collectively enable sensitivity across a broad spectrum of ultraviolet and visible light (Musilova et al., 2021). Visual sensitivity is further influenced by the binding of opsin proteins to either vitamin A1- or A2-derived retinal chromophores, which modulates the spectral sensitivity of opsins (Carleton et al., 2020; Musilova et al., 2021). Visual plasticity, allowing for adaptation to variable light conditions, is achieved through two primary, reversible mechanisms: modulation of opsin gene expression and the adjustment of chromophore composition (Carleton et al., 2020; Musilova et al., 2021). Species with multiple cone opsins can dynamically alter their relative expression levels in response to prevailing spectral conditions, a phenomenon observed in *Metriaclima mbenji*, which rapidly downregulates UV-sensitive SWS1 opsin when exposed to UV-deficient environments (Nandamuri et al., 2017). Similarly, exposure to turbid, red-shifted light induced upregulation of LWS opsin expression in the adult red shiner (*Cyprinella lutrensis*) (Chang & Yan, 2019). Furthermore, chromophore switching is enzymatically regulated, often by Cyp27c1; for instance, increased expression of Cyp27c1 facilitates the conversion of A1 precursors to A2, shifting peak absorbance toward longer wavelengths, whereas A1-dominance typically promotes shorter-wavelength sensitivity (Escobar-Camacho et al., 2019). Together, these mechanisms provide a flexible framework for fishes to optimize their visual sensitivity to match their specific environmental light regimes. While both mechanisms contribute to environmental adaptation, plasticity in opsin gene expression generally produces larger spectral shifts than A1/A2 chromophore exchange (Corbo, 2021; Carleton and Yourick, 2020).

Although numerous studies have explored the breadth and extent of visual plasticity in fishes, relatively few have investigated the functional significance of opsin expression plasticity, particularly in terms of altered visual perception, behavioural performance, and its potential consequences for fitness (Carleton et al., 2020; Bertinetti & Torres-Dowdall, 2025; Musilova et al., 2021; Carleton & Yourick, 2020). Among the studies that have examined its functional consequences, visual performance has been assessed using a range of behavioural tasks, from relatively simple reflexive responses, such as optomotor and dorsal light responses, to more complex behaviours involving responses to visual stimuli, foraging preferences, and mate choice (Fuller et al., 2010; Smith et al., 2011; Sakai et al., 2018; Wright et al., 2020; Potkamp et al., 2026). One such study by Sakai et al. (2018) demonstrated that preferences of female guppies (*Poecilia reticulata*) towards male colourations vary among populations and even across individuals within a population, and such variations were partially explained by the opsins expression patterns. Although these findings highlight the potential functional consequences of opsin expression plasticity for visually mediated social preferences, such integrative investigations have largely focused on a limited array of model organisms, including guppies, killifish, and certain cichlid lineages, predominantly the African Pseudocrenilabrinae. The ancient and phylogenetically distinct Etroplinae lineage remains largely unexplored, presenting an important gap in our understanding of visual plasticity within the broader cichlid radiation. To address this, we utilized the green chromide (*Etroplus suratensis*), a species inhabiting dynamic aquatic environments characterized by substantial spectral variability (Haque et al., 2024), as a model organism. Given that green chromide relies heavily on vision for conspecific recognition and foraging (XXX), demonstrably linked to the fitness of the species (XXX), we investigated plasticity of its socio-sensory network. By assessing the interplay between visual mechanisms and social preference across contrasting light environments, this study elucidates how green chromide maintains behavioral efficacy amidst fluctuating environmental conditions.

To investigate this, we initially characterized the visual pigments of the green chromide by identifying its retinal opsin genes. Given the absence of prior sequence data for this species, we employed RNA sequencing to delineate these genes, verifying their orthology to established vertebrate opsin classes via phylogenetic comparison with other cichlid sequences. Our analyses indicated a long-wavelength-biased visual sensitivity, characterized by the predominant expression of LWS and RH2A cone opsins under standard (white) light conditions. Leveraging this spectral baseline, we utilized blue-shifted light, a spectrum naturally encountered in clear, shallow aquatic habitats, as a contrasting experimental treatment. We subsequently evaluated shifts in visual physiological mechanisms, specifically opsin gene expression and chromophore composition, alongside behavioral performance in social preferences and foraging assays under both white and blue light regimes. These data suggest that the integration of visual sensory modulation and behavioral flexibility likely facilitates adaptive functionality in green chromides across heterogeneous light environments.

## 2 Methods

### 2.1 Study species

Green chromides were captured using cast nets and collected from the Hooghly River near Mousuni Island, West Bengal, India (21°41′44.88″ N, 88°11′8.714″ E), and tilapia (*Oreochromis* sp.) were caught using cast net from Petli lake, Kheda, Gujarat, India (22°35′56.861″ N, 72°45′26.604″ E) during December 2024-January 2025. The fish were transported to laboratory facilities under controlled conditions to minimise transport-related stress. During transportation, water temperature was maintained at approximately 25–28°C and dissolved oxygen was maintained above 5 mg L⁻¹ through continuous aeration. The journey lasted approximately 24 hours, no water changes required during transportation.

Individuals were housed in 70-L glass tanks and fed once daily (5% of their body weight) with commercially available freeze-dried bloodworms (Kishen Industries, Pune, India). The water temperature was maintained at 27 ± 2 °C and pH was maintained between 7 to 8, with a 12 h light:12 h dark photoperiod. Fish were acclimatised in the laboratory for at least six months before the experiments began. All the green chromides used in this study had a total body length of 5-8 cm (measured from the tip of the snout to the tip of the caudal fin; Onsoy et al., 2011), corresponding to a sexually immature stage (Vikas et al., 2020; Ward and Wyman, 1977). Tilapia used as heterospecific stimulus fish were size-matched to the focal green chromides (5–8 cm total body length). All individuals were visually screened to ensure they were free from external parasites and fin rot, and only healthy fish were used in the experiments.

### 2.2 Light environments

Green chromides were divided into two treatment groups. The first treatment group was housed for 90 days under two sequential light environments: (1) white light from day 0 (W0) to day 45 (W45), followed by (2) blue light from day 46 (B0) to day 90 (B45) (Figure 1a). The second group was maintained under white light for the entire 90-day period (W90). This allowed us to distinguish the effect of light environment from that of housing time. All the chromides were housed in 70 L glass tanks (53x37.5x38.5 cm, length x width x height) placed inside opaque black cardboard boxes (white-light environment: 62x43x64 cm; blue-light environment: 54x40x80 cm, length x width x height) and illuminated by a single LED bulb from above. Water temperature (27±2°C), photoperiod (12:12, light:dark), and feeding regime (commercial freeze-dried worm once a day at 5% of body weight) were kept identical across all housing tanks for each light environment. Care was taken to ensure that no outside light entered the boxes. Illumination was provided using a white LED bulb (9 Watt, Philips, India) for the first 45 days (and all throughout 90 days for the second group), followed by a blue LED bulb (9 Watt, Halonix, India) for the next 45 days. The absolute irradiance of the downwelling light (downward incident light from the source, bulb, towards the water surface) was measured, from 300 nm to 800 nm, in the housing tanks for each of the light environment, using a 230 µm optic fibre (UV-Vis XSR, Ocean Optics, USA) and a cosine corrector (CC 3-UVS, Ocean Insight, USA) connected to Avantes AvaSpec-ULS3648 spectrometer, Netherlands (Figure 1b) (Luehrmann et al., 2018). Equal irradiance across treatments was maintained by adjusting the bulb height to 64 cm for white light and 80 cm for blue light.

**Figure 1.**
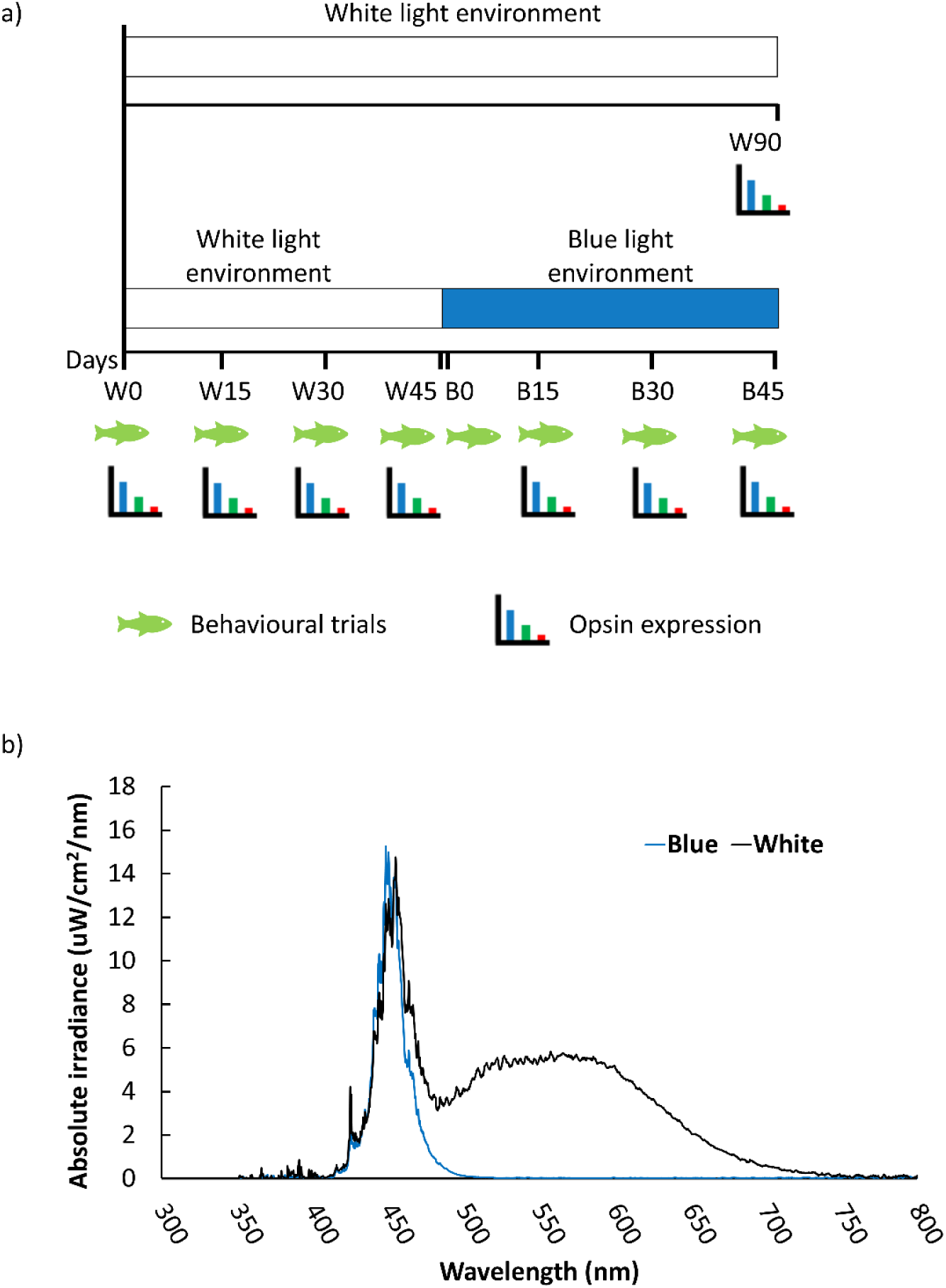
Light environment and experimental timeline for green chromides. a) Absolute irradiance spectra of downwelling light measured in housing and experimental tanks under white and blue lights. (b) Experimental timeline showing sequential exposure of green chromides to white light [Day 0 to 45 (W0–W45)] followed by blue light [Day 46 to 90 (B0–B45)]. A separate group of fish was maintained under white light for 90 days (W90). Behavioural trials (social preferences) and sampling for opsin expression were conducted at an interval of every 15 days throughout the 90-day experimental period, except for the W90 group, which was sampled only for opsin expression.

### 2.3 Experimental timeline and sample size

For assessing behavioural traits in the first treatment group, 10 individuals were housed in a single tank. The same set of fish was carried forward from the white-light phase to the blue-light phase to ensure consistent visual experiences in behavioral trials across light environments. The same 10 individuals (N = 80 trials; 10 individuals per time point) were repeatedly tested for behavioural assays at W0, W15, W30, W45, B0, B15, B30, and B45. The W45 and B0 behavioral trials were conducted 24 h apart. For both the zero-day (W0 and B0) trials, fish were first tested (for behavioural assays) under the respective light environments, before being transferred to their corresponding environments. For opsin expression analyses in the first treatment group, four individuals were sampled at day 0 (W0), before being introduced into the experimental housing tanks. The remaining experimental fish (N = 24) were housed in three tanks, with eight individuals per tank. Subsequently, four individuals were sampled at each time point: days 15 (W15), 30 (W30), and 45 (W45) under white light, followed by days 15 (B15), 30 (B30), and 45 (B45) under blue light, thus, adding up to a sample size of 28 for gene expression analyses.

For the second treatment group, a separate tank with a group of four individuals (the second group) was maintained continuously under white light until day 90 (W90) to distinguish changes associated with experimental duration from those associated with the light environment (Figure 1a), and were analysed for gene expressions at W90. Apart from the treatment groups, an additional tank containing eight fish was maintained under the conditions similar to the first treatment group, potentially serving as a backup in case of substitutions required due to mortality.

### 2.4 Pipeline for opsin analysis

As opsin expression data was not available for the green chromides, we sequenced retinal transcriptome of a green chromide for the white light day-0 timepoint. Tissue from both eyes was pooled and homogenized, and the homogenate was subsequently divided into two samples that were processed as technical replicates for transcriptome sequencing. Opsin genes were identified by annotating assembled sequences, which were confirmed through phylogenetic analysis. Using the identified sequences, we designed primers targeting opsin genes of green chromide, and subsequently measured opsin gene expression via quantitative real-time polymerase chain reaction (qRT-PCR). Below is the detailed methodology for each step in the pipeline.

#### 2.4.1 Transcriptome sequencing (RNA-seq) and analysis

RNA quality was checked using Agilent RNA TapeStation (Agilent Technologies, California, USA). Subsequently, the RNA-seq libraries were prepared from 200 ng of total RNA using the NEBNext® Ultra™ II Directional RNA Library Prep Kit (New England BioLabs) at Genotypic Technology Pvt. Ltd., Bangalore, India. The libraries were prepared with USER-mediated second-strand excision for strand specificity, ligated to Illumina-compatible adapters, enriched by 11 cycles of PCR, purified with NEBNext beads, quantified using a Qubit fluorometer, and assessed for fragment size distribution on an Agilent 4150 TapeStation system. Retinal RNA-seq libraries were sequenced on an Illumina NovaSeq X Plus platform using 150 × 2 paired-end chemistry. An average of 24.49 million reads (∼7.35 Gbp of paired-end sequence data) was generated per sample. Adapter sequences and low-quality reads were removed using Trim Galore, retaining reads with a Phred quality score >30 and a minimum read length of 20 bp. Following filtering, an average of 24.20 million reads (∼7.26 Gbp of paired-end sequence data) per sample was retained, corresponding to 98.8% of the raw reads. The retained reads were assembled de novo using rnaSPAdes v3.15.5 (Bushmanova et al., 2019; Luehrmann et al., 2018; Delroisse et al., 2016).

Basic assembly metrics (N50, average, maximum and minimum contig lengths) were calculated. Assembled transcripts were clustered with CD-HIT-EST (95% similarity) (Li &Godzik, 2006) to reduce redundancy. Coding regions were predicted using TransDecoder (Haas, BJ. https://github.com/TransDecoder/TransDecoder). The predicted coding sequences were annotated against all available Actinopterygii protein sequences in the UniProt database using BLAST (Altschul et al., 1990), and sequences showing >30% identity were retained for further analysis. A targeted annotation against Cichlidae protein sequences was subsequently performed using the same approach as described above to identify opsin genes, Cyp27c1 (gene responsible for A1 to A2 retinal conversion) and three housekeeping genes, EF1α (Elongation Factor 1-alpha), UBE2Z (Ubiquitin conjugating Enzyme E2 Z), and GAPDH (Glyceraldehyde-3-Phosphate Dehydrogenase).

Additionally we performed RNA-seq–guided de-novo genome annotation of green chromide to generate a genome-wide profile of predicted genes. Subsequently, the genes of particular interest, such as the visual opsin genes and Cyp27c1, were screened within our newly generated annotation file. The reference genome assembly (NCBI accession GCA_041004005.1) of green chromide was annotated using the BRAKER3 pipeline (Gabriel et al., 2024), which integrated extrinsic RNA-seq data from the fish eye tissue with homologous protein information to improve gene prediction accuracy. Protein sequences from Nile tilapia (*Oreochromis niloticus*), a cichlid species, were supplied as evidence to guide gene prediction. BRAKER3 predicted 59,842 predicted coding sequences (CDS) and corresponding amino acid (AA) translations, amongst which Cyp27c1 and six visual opsins were identified: LWS, RH1, RH2A, SWS1, SWS2A & SWS2B. These candidate gene models were further evaluated by BLAST similarity searches against the Nile tilapia sequences at both the nucleotide CDS and predicted amino acid levels. Sequences showing >90% sequence identity and >99% query coverage were retained as orthologous candidates. This ensured accurate identification and excluded paralogous or mis-annotated genes.

#### 2.4.2 Phylogenetic tree construction

Reference opsin sequences were systematically retrieved from the NCBI Nucleotide database using custom Python scripts. Searches were restricted to Cichlidae and performed independently for each target opsin using gene-specific NCBI annotations (RH1, SWS1, SWS2A, SWS2B, RH2A, and LWS). Species with sequence records for all five opsins (RH1, SWS1, SWS2A, SWS2B, and LWS) were initially retained. Species with RH2A sequence record were retained separately as there were only a few species, whose RH2A sequence was available. When multiple sequences were available for a given opsin within a species, the longest sequence was selected for downstream analyses. The corresponding GenBank accession numbers are provided in Table 1. For each opsin gene, the predicted amino acid sequence of green chromides was combined with orthologous sequences from other cichlid species (Table 1). Individual gene datasets were prepared for downstream alignment and phylogenetic inference. Protein sequences were aligned using the Conda-based standalone command-line tool MAFFT v7.525 (Katoh and Standley, 2013), with the L-INS-i iterative refinement strategy, which is suitable for divergent homologous proteins. To preserve codon structure for evolutionary analysis, the amino acid alignment was back-translated into codon alignment using the command-line-based tool PAL2NAL (Suyama et al., 2006) (see Data 1 for amino acid alignment files). The best-fit substitution model for each gene alignment was determined using ModelTest-NG (Darriba et al., 2019) under the Akaike Information Criterion (AIC). The selected model for each opsin dataset was then used in downstream maximum-likelihood phylogenetic tree reconstruction.

**Table 1:** List of species used for phylogenetic tree construction.

| Species |  | GenBank accession numbers |  |  |  |  |  |
|---|---|---|---|---|---|---|---|
| Scientific name | Common name | Genes |  |  |  |  |  |
|  |  | SWS1 | SWS2A | SWS2B | LWS | RH1 | RH2A |
| *Acarichthys heckelii* | Threadfin acara | MW401034.1 | MW588222.1 | MW591510.1 | MW588246.1 | MF078392.1 | MW588234.1 |
| *Amatitlania siquia* | Convict cichlid | MW591516.1 | MW591507.1 | MW591509.1 | MW588258.1 | KP715367.1 | MW588262.1 |
| *Aulonocara hueseri* | Night aulonocara | AY775100.1 | AY775074.1 | AY775083.1 | AY780517.1 | AY775112.1 |  |
| *Cynotilapia afra* | Afra cichlid | AY775104.1 | AY775079.1 | AY775088.1 | AY780521.1 | AY775118.1 |  |
| *Dimidiochromis compressiceps* | Malawi eyebiter | AF247125.1 | AF247113.1 | AF247117.1 | AF247125.1 | AY775059.1 | AF247130.1 |
| *Haplochromis degeni* | Degeni cichlid | AB666561.1 | AB667135.1 | AB666698.1 | AB667392.1 | AB667580.1 | AB667763.1 |
| *Haplochromis fischeri* |  | AB666574.1 | AB667150.1 | AB666711.1 | AB326136.1 | AB667604.1 | AB667770.1 |
| *Haplochromis pyrrhocephalus* |  | AB666550.1 | AB667127.1 | AB667125.1 | AB667370.1 | AB667558.1 | AB667740.1 |
| *Haplochromis macula* |  | AB666524.1 | AB667117.1 | AB666672.1 | AB667330.1 | AB667518.1 | AB667663.1 |
| *Labeotropheus fuelleborni* | Blue mbuna | AF191223.1 | AF247115.1 | AF247119.1 | AF247127.1 | AY775113.1 | AF247123.1 |
| *Lethrinops parvidens* | Lethrinops red flush | AY775102.1 | AY775077.1 | AY775087.1 | AY780519.1 | AY775116.1 |  |
| *Lucania goodei* | Bluefin killifish | AY296735 | AY296737 | AY296736 | AY296740 | AY296738 | AY296739.1 |
| *Maylandia zebra* | Zebra mbuna | NM001310074 | AF247114 | AF247118 | AF247126 | AY775114 | JF262089.1 |
| *Melanochromis auratus* | Golden mbuna | AY775101.1 | AY775076.1 | AY775084.1 | AY780518.1 | AY775115.1 |  |
| *Mylochromis lateristriga* | Basket hap | AY775105.1 | AY775075.1 | AY775085.1 | AY780522.1 | AY775119.1 |  |
| *Ophthalmotilapia ventralis* |  | AY775097.1 | AY775060.1 | AY775062.1 | AY780512.1 | AY775109.1 |  |
| *Oreochromis niloticus* | Nile tilapia | JF262087.1 | JF262088 | JF262088 | JF262088 | AY775108 | DQ235682 |
| *Oryzias latipes* | Japanese rice fish | AB223058.1 | AB223056.1 | AB223057.1 | AB435812.1 | AB180742.1 | AB223053.1 |
| *Pseudotropheus sp. acei* | Yellow-tail acei | DQ088642.1 | DQ088636.1 | DQ088639.1 | DQ088627.1 | KP130841.1 |  |
| *Pterophyllum scalare* | Freshwater angelfish | KX382945.1 | KX382946.1 | KX382955.1 | KX382917.1 | MF078457.1 | MW588241.1 |
| *Pundamilia nyererei* |  | AY673728.1 | AY673763.1 | AY673708.1 | AB448595.1 | KP130837.1 |  |
| *Stigmatochromis modestus* |  | AY775107.1 | AY775080.1 | AY775066.1 | AY780523.1 | AY775122.1 |  |
| *Tyrannochromis maculatus* |  | AY775103.1 | AY775078.1 | AY775086.1 | AY780520.1 | AY775117.1 |  |

Phylogenetic relationships were inferred using the maximum-likelihood framework implemented in IQ-TREE 2 (Minh et al., 2020). Branch support was assessed using 1,000 ultrafast bootstrap replicates, with 1,000 iterations applied to the ultrafast bootstrap convergence criterion to ensure stability of support values. To avoid local optima and ensure recovery of the global maximum-likelihood topology, ten independent tree searches were performed, and the tree with the highest log-likelihood score was selected as the final topology (Nguyen et al., 2015; Minh et al., 2020). Convergence was assessed by confirming repeated recovery of the optimal likelihood across independent runs. Two non-cichlids, Bluefin killifish (*Lucania goodei*), belonging to the family Fundulidae, and Japanese rice fish (*Oryzias latipes*), belonging to Adrianichthyidae, were used as an outgroup to root all phylogenetic trees, allowing directional inference of opsin evolution within cichlids. A comprehensive multi-gene phylogenetic analysis was also performed. The six individual curated datasets (LWS, SWS1, SWS2A, SWS2B, RH1, and RH2) were merged into a single composite dataset containing all 131 sequences from 25 species (23 species for SWS1, SWS2A, SWS2B, and LWS; 14 species for RH2A) (Table 1) (see Data 2 for tree files). Phylogenetic reconstruction of this super-alignment was performed in IQ-TREE 2, where the best-fit amino acid substitution model was automatically selected using ModelFinder Plus (MFP) based on the Bayesian Information Criterion (BIC) (Minh et al., 2020). Branch support was estimated using 1,000 ultrafast bootstrap replicates.

The final trees were visualized and annotated using iTOL v7.4.2 (Interactive Tree of Life) (Letunic & Bork, 2024), with branch support values used to assess support for the inferred clades. Phylogenetic analysis was omitted for Cyp27c1 because functional adaptation in this enzyme is driven by differential gene expression rather than coding-sequence mutations or single nucleotide polymorphisms (SNPs), and orthology was confirmed by high sequence identity (>85% identity) across available Cichlidae orthologs (see data 3 for Cyp27c1 sequence alignments).

#### 2.4.3 qRT-PCR standardization

Primers were designed for all the identified genes (Table S1) using Primer3 (Untergasser et al., 2012) software, based on conserved regions having orthologous sequences from Nile tilapia (*Oreochromis niloticus*), zebra mbuna (*Metriaclima zebra*), blue mbuna (*Labeotropheus fuelleborni*), and Malawi eye biter (*Dimidiochromis compressiceps*). Primer design parameters were set to a GC content of 40–60% and a melting temperature (T_m_) of 58–62°C. The specificity of each primer pair was verified *in silico* using NCBI Primer-BLAST, and Integrated DNA Technologies (IDT) was used to check for the primer-dimers and hairpin loops. PCR amplification was performed, and the amplified products were visualized on an agarose gel to confirm the expected product sizes. PCR efficiency for each primer pair was determined using four-point (50, 25, 12.5, and 6.25 ng per reaction) serial dilutions of cDNA. The efficiency (E) of each reaction was calculated from the slope of the standard curve using the formula (Ward et al., 2008):

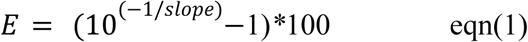

Efficiencies between 85–105% were considered acceptable (Carleton and Kocher, 2001; Stieb et al., 2016; Fuller et al., 2004). Efficiency values and primer sequences for each gene are provided in Table S1.

#### 2.4.4 Standardization of qPCR assays and quantification of opsin expression

Green chromides were euthanized using an overdose of clove oil. Euthanasia was performed at a fixed time, between 11:00 a.m. and 1:00 p.m., across all the sampling intervals to minimize variation due to diurnal effects (Yourick et al., 2019). Both eyes were carefully dissected from each individual, and the tissues were immediately preserved in RNAlater (Invitrogen, Carlsbad, CA, USA) to preserve RNA integrity (Yourick et al., 2019). RNA was extracted on the same day. During RNA extraction, both the eyes from each fish were pooled and homogenized together using liquid nitrogen to get a representative sample of the individual’s eye tissue (Zhang et al., 2022). Total RNA was extracted from the pooled tissue using the TRIzol method followed by purification using the RNeasy Micro Kit (Qiagen, Hilden, Germany) (Zhang et al., 2022). RNA quality and quantity were determined using the DeNovix DS-11 spectrophotometer (DeNovix Inc., Wilmington, DE, USA). A total of 1000 ng of high-quality RNA from each sample was reverse-transcribed to complementary DNA (cDNA) using the iScript cDNA Synthesis Kit (Bio-Rad, Hercules, CA, USA) (Luehrmann et al., 2018).

qPCR (Quantitative PCR) was performed using the KAPA SYBR FAST qPCR Master Mix (Roche, Basel, Switzerland) on the QuantStudio 3 Real-Time PCR System (Applied Biosystems, Foster City, CA, USA). The 20 µL reaction included 10 µL of 2× KAPA SYBR mix, 0.5 µL forward and reverse primers (10 µM) each, and 15 ng of cDNA template, with nuclease-free water added to complete the volume. The qPCR cycling conditions were as follows: the first denaturation at 95 °C for 3 min was followed by 40 cycles of 95 °C for 3 s, and 60°C for 20 s (annealing and extension). At the end of each run, melt-curve analysis was performed to confirm amplification specificity. All samples were analysed twice on different plates for opsins and once for Cyp27c1, with each run performed in triplicate. The three Ct values from each plate were averaged, and the resulting mean Ct values were normalised. The normalised opsin values from the two plates were then averaged and used for further analysis (Luehrmann et al., 2018). We performed two types of normalisation on Ct values: independent (relative to the housekeeping gene) and dependent (expressing each cone opsin as a percentage of the total expression of the five cone opsin genes). Independent normalisation allowed assessment of changes in transcript abundance relative to the housekeeping gene, whereas the dependent approach captured within-retinal shifts in spectral sensitivity by quantifying changes in the relative contribution of each opsin class within the cone mosaic. This becomes particularly important in systems where multiple opsins can be coexpressed within single photoreceptors and fine-scale adjustments in transcript abundance may shift the spectral phenotype (Nandamuri et al., 2017; Stieb et al., 2016). However, Cyp27c1 expression was normalised using only the independent normalisation method.

Independent opsin normalisation was performed using equation (2) (Yourick et al., 2019) .

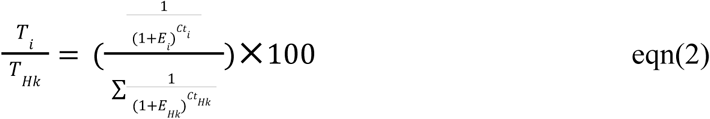

*T_i_/T_Hk_* is the expression of each individual opsin, i, relative to the expression of housekeeping, Hk. E_i_ is the PCR efficiency for each primer pair, and *Ct_i_* is the average critical cycle number for each opsin gene and, similarly, *Ct_HK_* for the housekeeping gene. Although all six opsin genes were measured with respect to a housekeeping gene, the expression of each opsin gene is independent of the expression of the other opsins. Independent normalisation was performed using a housekeeping gene that maintained stable (low variations) expression across the two light treatments. To identify this gene, the expression stability of all candidate housekeeping genes was evaluated using RefFinder (https://www.ciidirsinaloa.com.mx/RefFinder-master/), a comprehensive web-based tool that integrates four commonly used algorithms: geNorm, NormFinder, BestKeeper, and the comparative ΔCt method (Xie et al., 2012). Ct values of each housekeeping gene were analysed in RefFinder using these four approaches. The gene showing the highest overall expression stability (lowest ranking value or stability score) was selected for normalization of opsin gene expression (Yourick et al., 2019).

The dependent normalisation was performed according to equation (3) (Yourick et al., 2019)

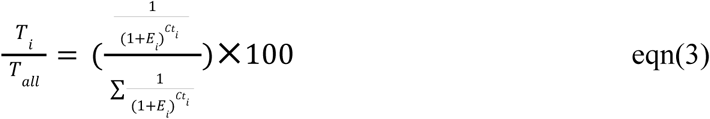

*T_i_/T_all_* is the proportional gene expression for a given gene (Luehrmann et al., 2018; Carleton and Kocher., 2001). When expressed as percentages, the five cone opsins together sum to 100% for each individual. The terms *E_i_* and *Ct_i_*are the same, as explained above.

### 2.5 Behavioural trials (social preferences)

To examine the functional significance of opsin expression under the given light environment, we conducted social preference trials in green chromides under similar light conditions (white and blue) as was maintained for housing the fish. All behavioural trials were conducted in a 100 L glass tank (36×40×70 cm, length × height × width), referred to as experimental tanks from here onwards. The experimental tank was divided into three chambers (left, right, and middle) using transparent, non-perforated plexiglass partitions (Hiral Enterprise, Ahmedabad, India) to prevent water exchange and thereby blocking olfactory cues (XXX). The middle chamber was further divided into three zones: left (L) and right (R) preference zones, and a central non-preference zone (C), each 10 cm wide (XXX). The light environment for the experimental tanks corresponded to that of the housing tanks and the irradiance profile was maintained using the same protocol as described above. Reaction to food was monitored in the housing tanks under both light conditions to examine whether the ambient light regime influenced foraging behaviour. Fish were fed every morning with commercially available freeze-dried bloodworms (Kishen Industries, Pune, India) at 5% of their body weight, provided as floating food, and their response to the food stimulus was recorded.

Behavioural trials (N = 80; 10 trials per time-point for each light treatment) to measure social preferences were conducted following the protocol as described in XXX. Each trial consisted of three consecutive phases, each of 10 minute duration and separated by 1-minute of acclimatisation: phase 1: 0 vs 0 (no stimulus on either side; negative control), phase 2: 0 versus green chromide (two green chromides in one of the side chambers), and phase 3: green chromide versus tilapia (two green chromides in one side chamber and two tilapia in the other). A single test fish (green chromide) was placed in the central chamber and was given one minute to acclimatise before the trial began. Stimulus fish were added at the start of their respective phases, and the stimulus and test fishes were size-matched. In each trial, the side (left or right) chamber in which the stimulus fish was placed, was randomised across trials to control for potential side-based biases. All the social preference trials were video recorded using a digital handycam (Panasonic HC-V385). Each test fish was used only once, and stimulus fish were randomly selected from holding tanks that were separately maintained for each species. All experiments were carried out under low-noise conditions. Social preferences were quantified using a relative preference index. The position of the test fish in the left (L) and right (R) zones of the middle chamber was recorded every 15 s during all the phases. Relative preference (RP) was calculated as OS/OT, (Durrer et al., 2020), where OS is the total number of occurrences of the test fish in the zone designated as the stimulus side. In all the phases (0 versus green chromide, and green chromide versus tilapia), the stimulus side always corresponded to the side where the stimulus green chromide (GC) was introduced. Since 0 versus 0 trials were subsequently followed by the 0 versus green chromide trials, we chose the side of the stimulus green chromides as the stimulus side for 0 versus 0 trials, as well. OT is the total number of occurrences of the test fish in both the left (L) and right (R) zones of the middle chamber. RP was calculated separately for all the three phases of the behavioural trials.

### 2.6 Statistical analysis

For independent opsin and Cyp27c1 expression, data was first tested for normality using the Shapiro–Wilk test and for homogeneity of variances using Levene’s test. Depending on these results, either one-way ANOVA or the Kruskal Wallis test was performed across time points (W0 to B45). When a significant difference was detected, post-hoc comparisons were performed using Dunnett’s test (for ANOVA) or Dunn’s test (for Kruskal–Wallis), between W45 (considered as endpoint for control, the white light conditions) and each of the blue-light time points (i.e., B15, B30 and B45).

To assess whether dependent opsin expression changed across the light treatments, we performed quasibinomial GLM (generalized link model) with a logit link, which is well-suited for modelling proportional data bounded between 0 and 1, and accounts for potential overdispersion in the data (Dunn & Smyth, 2018). For each opsin gene, we fitted an interaction-based quasibinomial model with a logit link function with opsin expression as the response variable and day (i.e., Day0, Day15, Day30, Day45) and light (white, blue) as categorical predictors (expression∼day*light). W45 was set as the reference (baseline) level for all models and for all the post-hoc comparisons. Post-hoc comparisons of W45 versus B15, B30 and B45, were conducted using the emmeans package in R, with Bonferroni correction applied to adjust for multiple testing.

For cone opsins showing significant effects in the above analyses, we additionally tested whether these changes (in the dependent opsin expression) were attributable to blue-light exposure or due to the housing conditions, by fitting a separate quasibinomial model with a logit link function (expression∼treatment) including W45, W90 and B45, with B45 as reference (baseline) point in the model. In this case, post-hoc comparisons were conducted among all the treatments using emmeans with Bonferroni correction.

Additionally, we also assessed the relationship between opsin expression and Cyp27c1 expression for each light condition by using Spearman’s rank correlation test. We used dependent-normalised expression data for opsin and independent-normalised expression data for Cyp27c1. Only the opsin genes that showed a significant day×light interaction in the GLM-based analysis were selected for this downstream analysis.

To determine whether social preferences changed under blue-light conditions when compared to white-light setup, preference values of the green chromide vs tilapia trials (bounded between 0 and 1) were modelled using quasibinomial GLM with preference as a response variable, and day and light as categorical predictors (preferences∼day*light) in the model. Post-hoc tests comparing W45 with B0, B15, B30, and B45 were conducted using emmeans with Bonferroni correction.

A significance threshold of α=0.05 was used for all data analysis. All analyses were performed in R (version 4.3.2).

## 3 Results

### 3.1 Identification of retinal opsins in green chromides

*De novo* transcript assembly reconstructed the complete coding sequences of six different opsin genes (rod opsin: RH1; cone opsins: SWS1, SWS2A, SWS2B, RH2A, and LWS), housekeeping genes (EF1α, UBE2Z, and GAPDH), and Cyp27c1 in green chromides. Details of the gene sequences are provided in Table S2 and the transcript values from RNA-seq for each opsin gene are given in Table S3. Identity of opsin transcripts was confirmed by amino acid-based maximum-likelihood phylogenetic analysis (figure 2).

**Figure 2.**
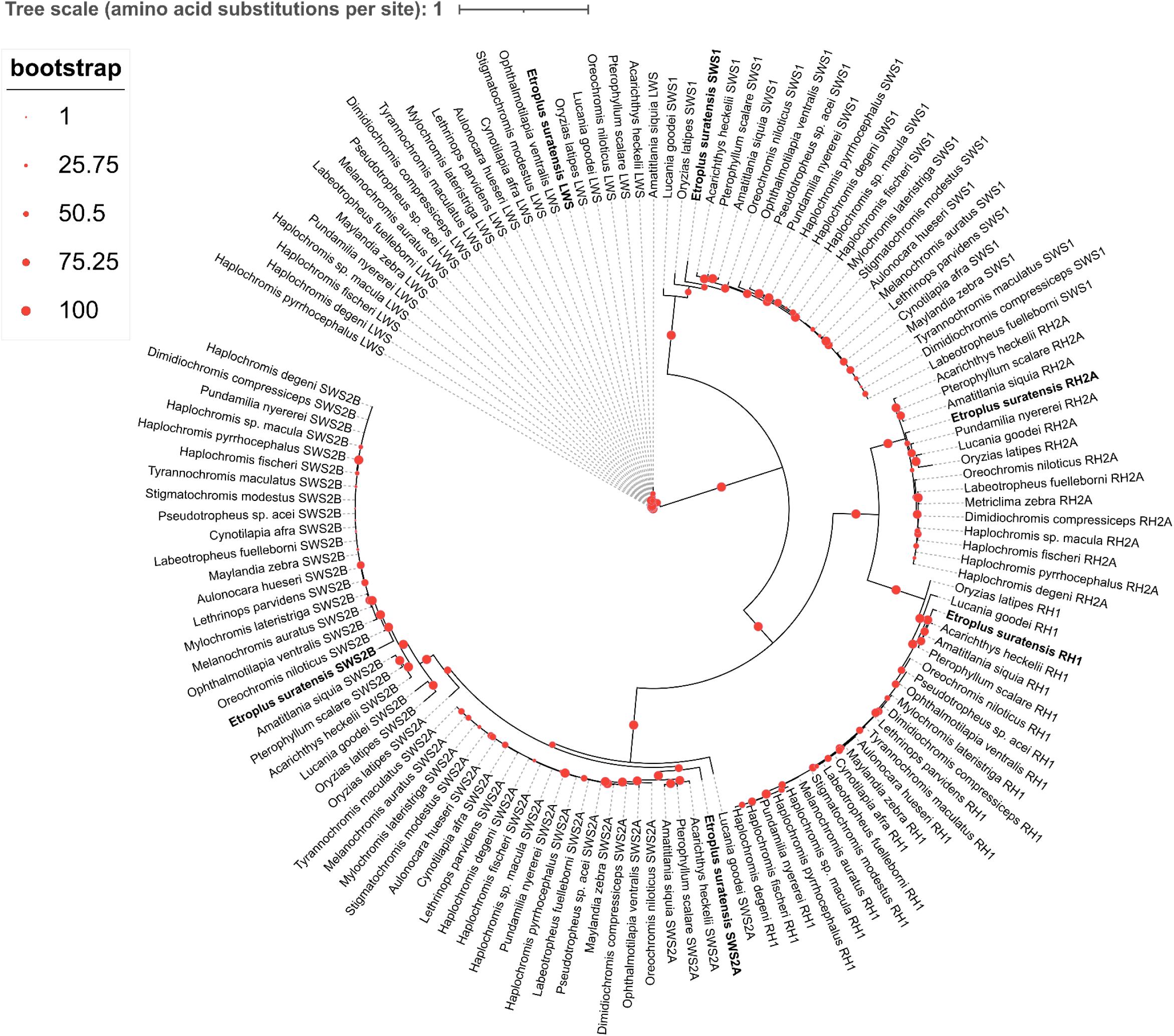
Maximum-likelihood (ML) phylogenetic tree of visual opsins in green chromides and other cichlids. Maximum-likelihood phylogeny based on amino acid sequences of six opsin genes (RH1, SWS1, SWS2A, SWS2B, RH2A, and LWS), showing the placement of green chromide opsins among cichlid visual opsins. Lucania goodei and Oryzias latipes were used as outgroups. Node circles indicate ultrafast bootstrap support values based on 1,000 replicates.

### 3.2 Identification of a stable housekeeping gene

Using RefFinder, the housekeeping genes were ranked from most to least stable, starting from UBE2Z (1.00), EF1α (1.68), and GAPDH (3.00), with stability scores shown in parentheses. Both UBE2Z and EF1α exhibited high stability across all four algorithms. However, UBE2Z showed relatively low expression levels, whereas EF1α was more abundantly expressed. Given that very low expression can reduce the reliability of normalisation, EF1α was selected as the reference gene for independent normalisation of opsin gene expression (Koreza and Rapacz, 2013).

### 3.3 Gene expression

#### 3.3.1 Independent normalisation of gene expression across time points

##### 3.3.1.1 Independent normalisation of opsin expression

For all the opsins, independent normalisation values did not show any significant differences across time points (One-way ANOVA: RH1, F=0.676, P=0.69; SWS2A F=1.043, P=0.429; SWS2B, F=0.291, P=0.951; RH2A, F=0.515, P=0.814 and LWS, F=0.552, P=0.787; Kruskal Wallis: SWS1, Χ^2^=2.849, P=0.899) (Figure 3). The full dataset spanning W0 to B45 is presented in Figure S1.

**Figure 3.**
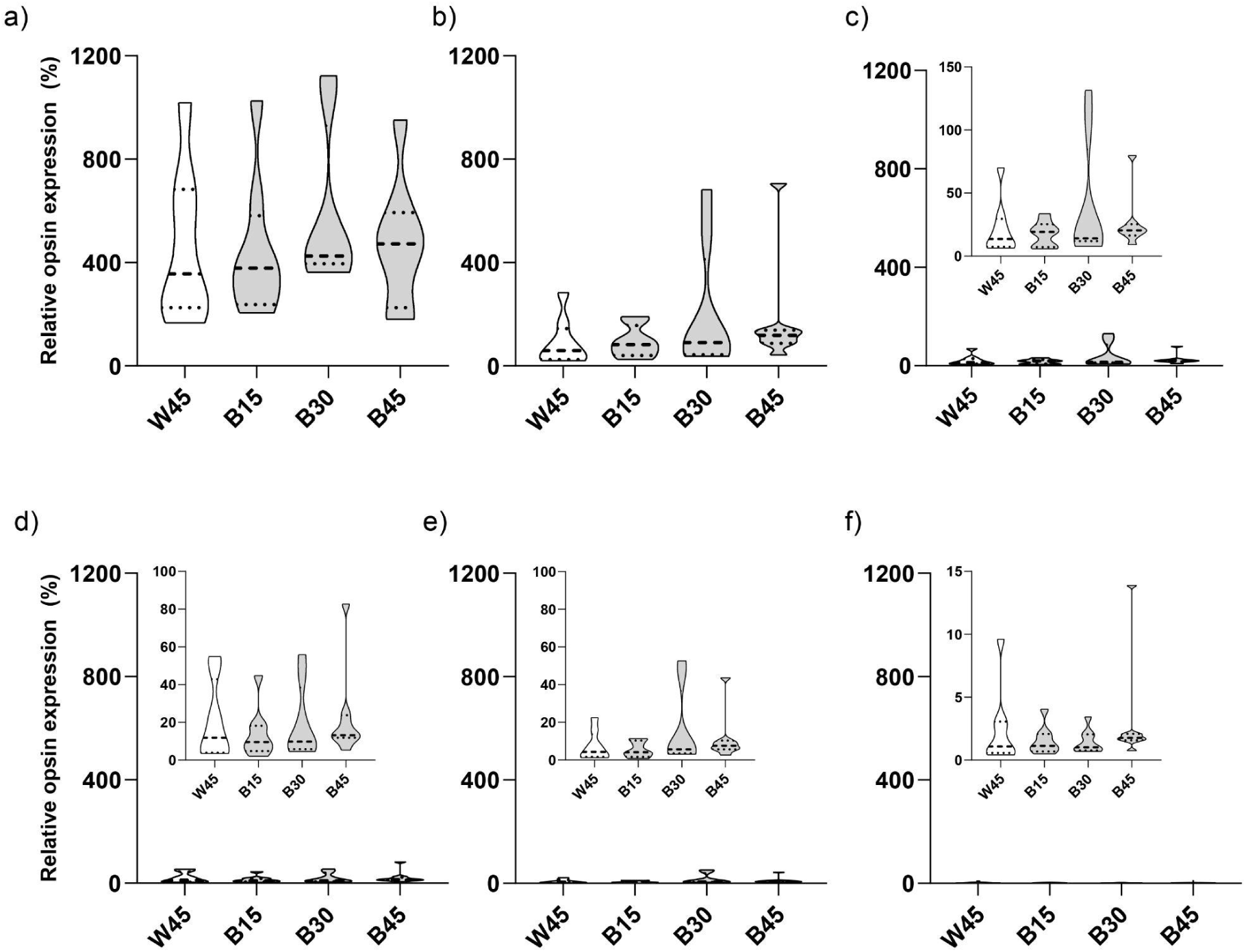
Independent normalisation of opsin expression. Violin plots illustrate relative expression (Ti/THk) of opsin genes, expressed as a percentage relative to EF1α. Panels show expression of (a) RH1, (b) LWS, (c) RH2A, (d) SWS1, (e) SWS2A, and (f) SWS2B genes. White and grey violins indicate white- and blue-light treatments, respectively. Each violin depicts the distribution of normalized expression values, with each individual contributing two data values (one from each of two independent plate runs). The dashed black line within each violin represents the median, while the dotted black lines represent the upper and lower quartiles. Insets within each panel present scaled representations of the same data to enhance data visualization.

##### 3.3.1.2 Independent normalisation of Cyp27c1 expression

Expression of Cyp27c1 was significantly different across time points (Kruskal Wallis: Χ^2^=14.231, P=0.027). Post-hoc tests revealed significant differences at B15 (P=0.04) and B45 (P=0.036) when compared to W45 (Figure 4). This suggests reduced Cyp27c1-mediated conversion of A1 to A2 under blue light, consistent with greater reliance on A1 chromophore. The full dataset spanning W0 to B45 is presented in Figure S2.

**Figure 4.**
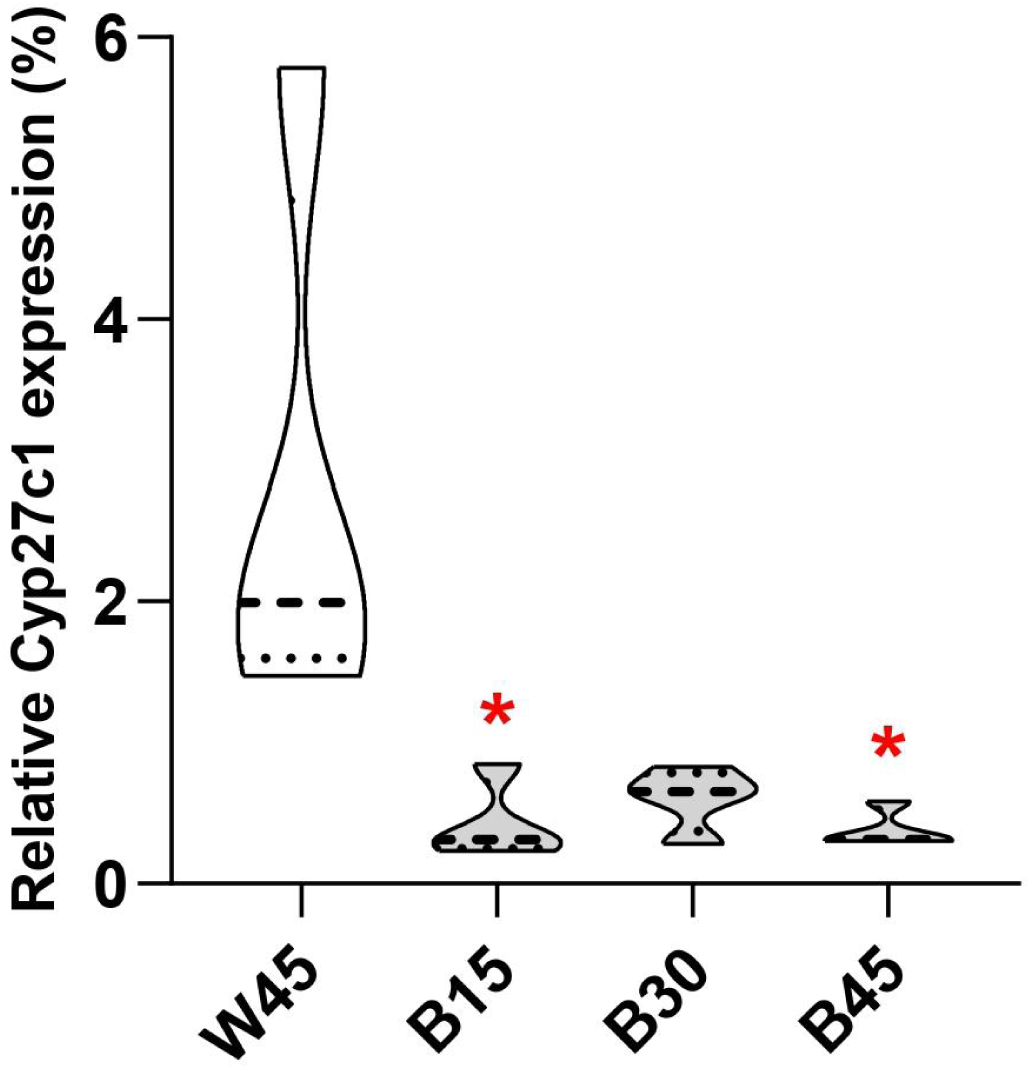
Independent normalisation of Cyp27c1 expression. Violin plots illustrate relative expression (Ti/THk) for Cyp27c1 gene, expressed as a percentage relative to EF1α. White and grey violins indicate white-and blue-light treatments, respectively. Each violin depicts the distribution of normalized expression values, with each individual contributing one data value (from a single plate run). The dashed black line within each violin represents the median, while the dotted black lines represent the upper and lower quartiles. An asterisk (*) indicates significant (Kruskal Wallis using post-hoc Dunn’s test) differences when compared to W45.

To test whether changes in Cyp27c1 expression were driven by blue-light exposure rather than an artefact of time, we compared Cyp27c1 expressions at W45, W90, and B45 treatments (Figure 5). The overall differences among treatments were significant (ANOVA: F = 16.15, P = 0.001). Post-hoc comparisons indicated that Cyp27c1 expression was significantly reduced under blue light (B45) relative to W45 and W90 (P = 0.001 and P = 0.019, respectively). No significant differences were detected between W45 and W90 (P = 1). Overall, these results indicate that the reduction in Cyp27c1 expression was associated with blue-light exposure rather than prolonged experimental duration.

**Figure 5.**
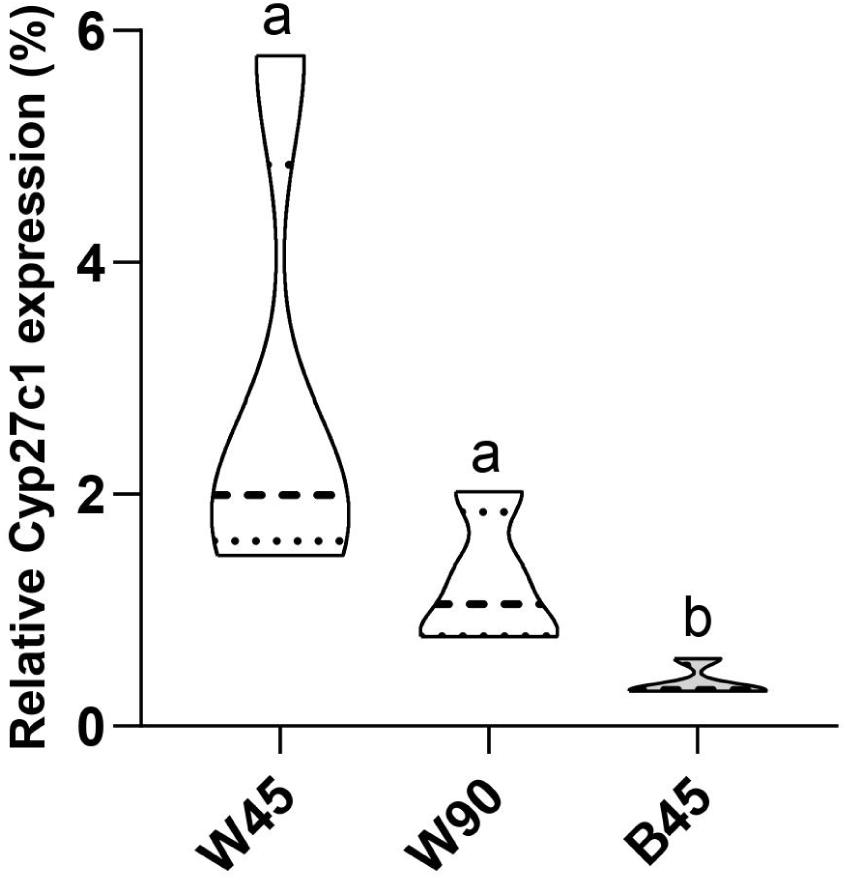
Independent normalisation of Cyp27c1 expression to test the effect of time. Violin plots illustrate relative Cyp27c1 expression across treatments, expressed as a percentage relative to EF1α. White and grey violins represent white- and blue-light conditions, respectively. Each violin depicts the distribution of normalized expression values, with each individual contributing one data value (from a single plate run). Dashed lines indicate median values, while dotted lines represent interquartile ranges. Differences among treatments were assessed using post-hoc Dunn’s test following a Kruskal–Wallis test, with significant differences indicated by different letters.

#### 3.3.2 Dependent normalisation of opsin expression across time points

Dependent normalisation revealed changes in relative cone opsin expression for LWS, RH2A, and SWS2A based on quasibinomial GLM analyses (Figure 6). The models showed significant main effects of day and light×day interactions, for LWS and RH2A (Table S4) and significant main effects of light and light×day interactions for SWS2A. SWS1 and SWS2B did not show a significant light×day interaction, although SWS2B expression showed a significant main effect of day (Figures 6c and 6d) (Table S4). For LWS, post-hoc comparisons indicated increased expression under blue light, with significant differences observed at B15 (P=0.009), B30 (P=0.047), and B45 (P=0.005) compared to W45 (Figure 6a). For RH2A, post-hoc analyses revealed a significant decrease in expression under blue light at B45 (P=0.021), whereas no significant differences were detected at earlier time points (B15, P=0.286; B30, P=0.588) when compared to W45 (Figure 6b). For SWS2A, post hoc pairwise comparisons did not reveal any significant differences (B15, P=0.446; B30, P=1; B45, P=1) relative to W45 (Figure 6e). The complete dataset of dependent normalization, spanning W0 to B45, for all the opsin gene expressions is presented in Figure S3.

**Figure 6.**
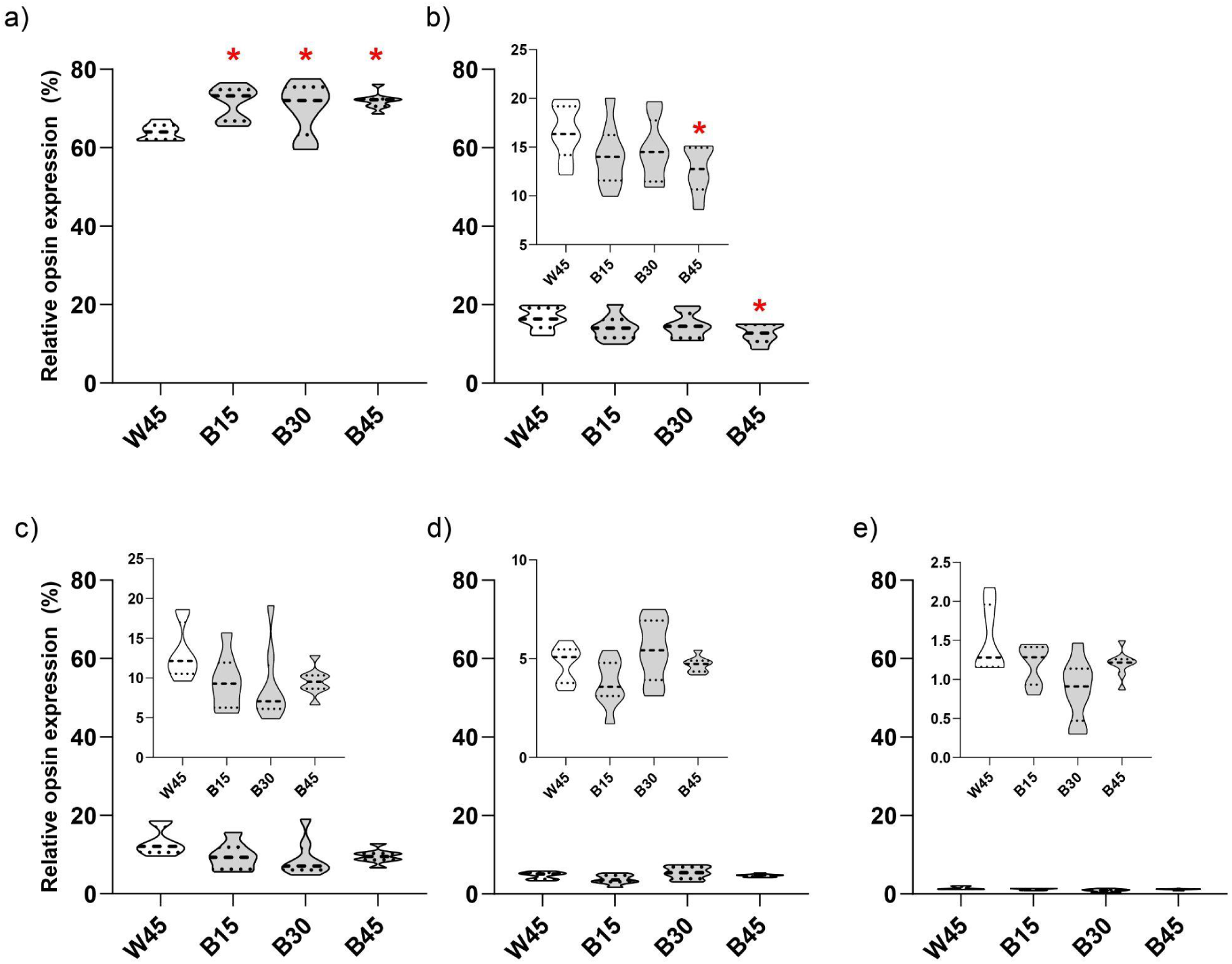
Dependent normalisation of cone opsin expression. Violin plots illustrate relative expression for each opsin gene expressed as a percentage. Panels show expression of (a) LWS, (b) RH2A, (c) SWS1, (d) SWS2A, and (e) SWS2B genes. White and grey violins indicate white- and blue-light treatments, respectively. Each violin depicts the distribution of normalized expression values, with each individual contributing two data values (one from each of two independent plate runs). The dashed black line within each violin represents the median, while the dotted black lines represent the upper and lower quartiles. Insets within each panel present scaled representations of the same data to enhance data visualization. An asterisk (*) indicates significant (quasibinomial GLM model using post-hoc comparisons) differences when compared to W45 (control).

To test whether changes in LWS and RH2A expression were driven by blue-light exposure rather than an artefact of time, we compared dependent opsin expressions at W45, W90, and B45 treatments (Figure 7). Quasibinomial GLM models for LWS and RH2A showed significant treatment effect (Table S5). Post-hoc tests revealed that LWS expression increased under blue light (B45) compared to W45 (P<0.0001) and W90 (P=0.0326), while W90 expression was not significantly different from W45 (P=0.064) (Figure 7a). RH2A expression decreased under blue light (B45) relative to W45 (P=0.006), while there were no significant differences between W45 and W90 (P=0.874), B45 and W90 (P=0.133) (Figure 7b). Overall, blue-light exposure primarily altered LWS expression, with no significant change in the expression of short-wavelength-sensitive opsins.

**Figure 7.**
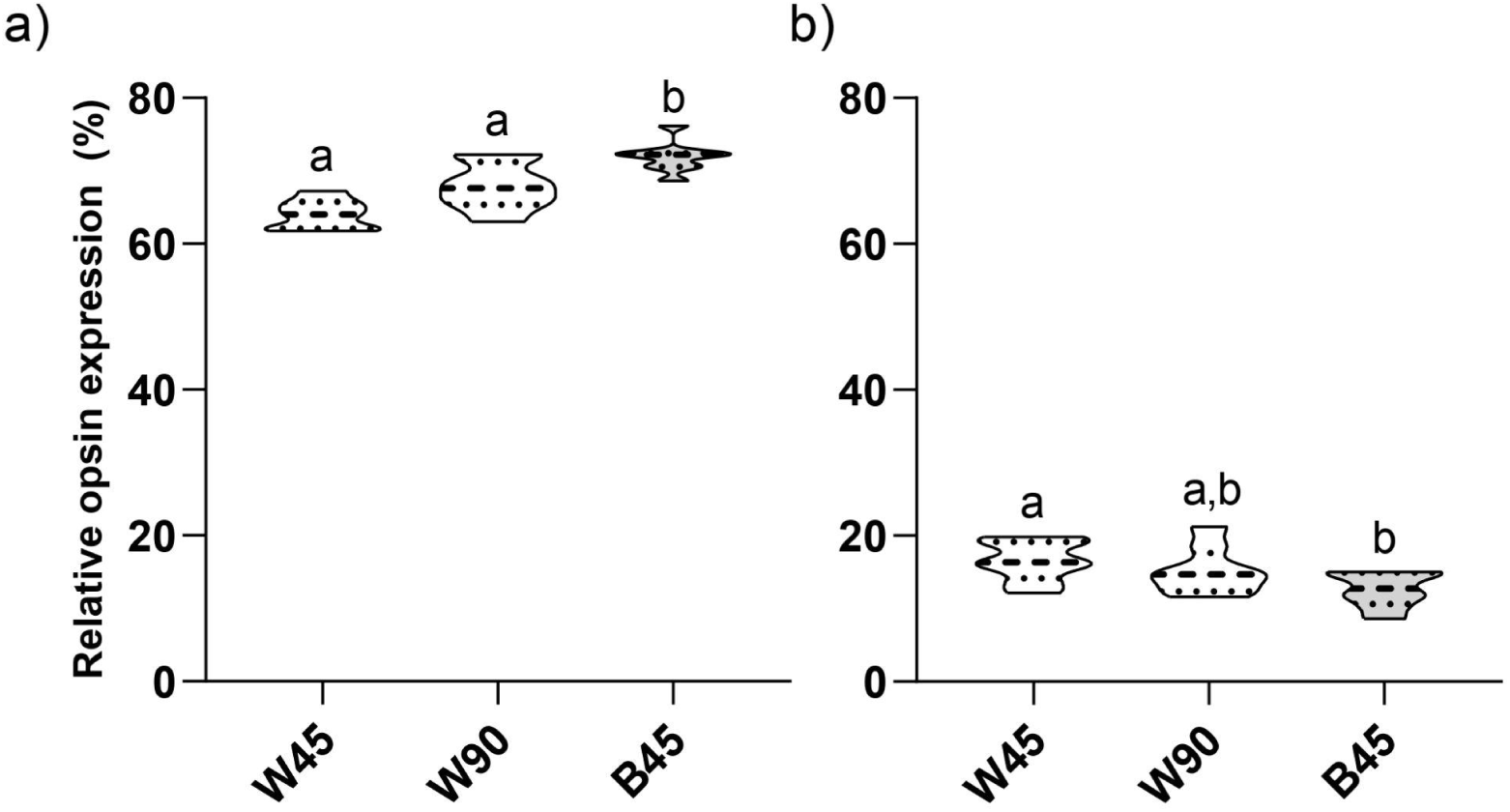
Dependent normalisation of LWS and RH2A expression to test the effect of time. Violin plots illustrate relative expression for each opsin gene expressed as a percentage. Panels show expression of (a) LWS, and (b) RH2A genes. White and grey violins indicate white- and blue-light treatments, respectively. Each violin depicts the distribution of normalized expression values, with each individual contributing two data values (one from each of two independent plate runs). The dashed black line within each violin represents the median, while the dotted black lines represent the upper and lower quartiles. Different alphabets indicate significant differences (using post-hoc pairwise comparisons in quasibinomial GLM models).

#### 3.3.3 Comparison of LWS and Cyp27c1 expression

There was no significant relationship between LWS expression and Cyp27c1 expression under white light (Spearman’s ρ = −0.31, n = 16, P = 0.23). Similarly, no significant relationship was detected under blue light (Spearman’s ρ = −0.18, n = 16, P = 0.52). White light showed greater variation compared to blue light, as indicated by the wider spread of the 95% confidence intervals (Figure S4).

### 3.4 Behavioural trials (social preferences)

Across all sampling intervals, green chromides consistently preferred conspecifics over tilapia, and this preference remained similar even under blue-light conditions, for all the different trial types including green chromide versus tilapia (Figure 8) and 0 versus green chromide (Figure S5). The quasibinomial GLM model (Table S6) on green chromide versus tilapia trials, revealed no significant effects of light, day, or their interaction. For the 0 versus green chromide trials (Figure S5), green chromides exhibited preference values of either 1 or close to 1, indicating a strong preference for conspecifics. 0 versus 0 trials (Figure S6), showed a random pattern under both the light conditions. Reaction to respond to food in the housing tanks was likewise comparable across the light treatments (Mean±SE, 5.0±2 seconds under white light and 4.8±2 seconds under blue light), indicating that foraging behaviour was unaffected by the ambient light environment. The full dataset spanning W0 to B45 for green chromide versus tilapia is presented in Figure S7.

**Figure 8.**
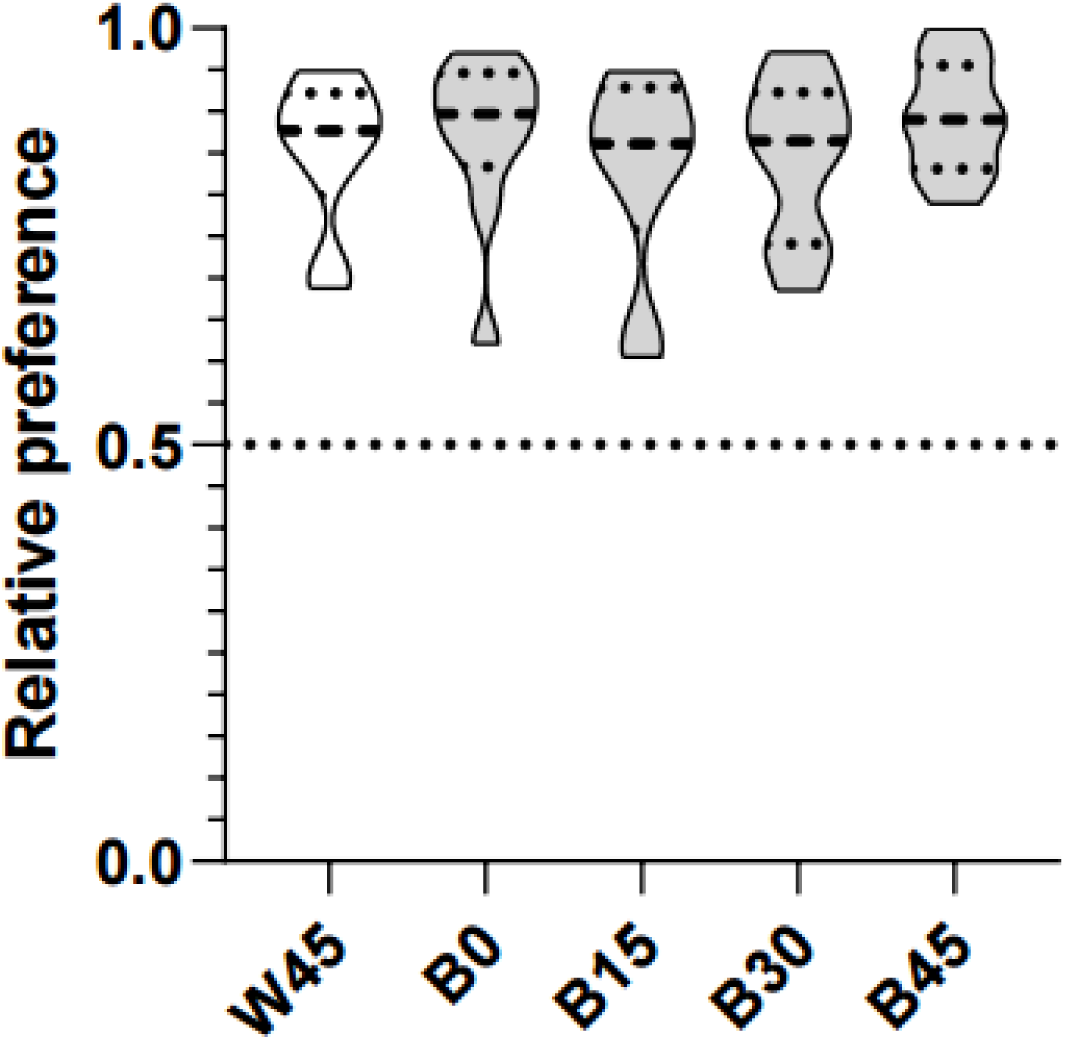
Social preferences of green chromides (green chromide versus tilapia trials) Violin plots illustrate social preferences of green chromides during green chromide versus tilapia trials. White and grey violins indicate white- and blue-light treatments, respectively. The dashed black line within each violin represents the median, while the dotted black lines represent the upper and lower quartiles. A dashed line at 0.5 relative preference indicates random preference

## 4 Discussion

Flexible visual processing enables organisms to navigate shifting light regimes, yet how molecular adjustments in the visual system interface with behavioral performance remains unclear. Here, we examined changes in visual-system components and social preferences in green chromides, exposed to different light environments. Under white-light conditions, green chromides exhibited a long-wavelength-biased opsin expression profile, with relatively high expression of LWS and RH2A. Dependent normalization (relative to total cone-opsin expression) revealed increased LWS and decreased RH2A expression under blue light, while short-wavelength-sensitive opsins remained largely unchanged. In contrast, independent normalization using a housekeeping gene revealed no significant differences in opsin expression across light conditions. This discrepancy between the two normalization approaches may arise because inter-individual variation in housekeeping gene expression can mask subtle changes in opsin transcript abundance, whereas dependent normalization captures changes in the relative contribution of individual opsins to the total cone-opsin pool (Bustin et al., 2009; Bustin et al., 2013). Cyp27c1 expression also decreased under blue light, consistent with reduced A1-to-A2 chromophore conversion and a greater relative contribution of the A1 chromophore. Parallel to these molecular shifts, green chromides maintained a consistent preference for conspecifics across all light conditions and showed similar feeding response times regardless of the spectral environment, with no significant lag in behavioral responses across contrasting light environments. Taken together, these results show that exposing the fish to a new spectral environment alters various molecular components of their visual machinery without causing any observable shifts in the social and foraging behaviors evaluated.

Increased LWS expression and decreased Cyp27c1 expression under blue light, when compared with W90 controls, confirmed that these shifts were driven by the spectral change rather than time-dependent effects. Furthermore, the down-regulation of Cyp27c1 indicates a shift toward the A1 chromophore under blue light, thereby tuning spectral sensitivity to shorter wavelengths, a pattern typical of teleosts in clearer photic regimes (Carleton et al., 2020; Bertinetti et al., 2024). Notably, LWS and Cyp27c1 expression levels were not significantly correlated under either light condition, mirroring the findings in Lake Victoria cichlids (Wilwert et al., 2022). This lack of co-regulation indicates that chromophore regulation and opsin transcription may operate via unlinked pathways. Consequently, while Cyp27c1 responded predictably to blue light, the opsin response departed from expectations: LWS expression increased from red to blue light treatments, while short-wavelength opsins (SWS1, SWS2A, and SWS2B) remained static. A similar comparable shift was observed in the turbid-water Neotropical cichlid, *Aequidens pulcher*, which developed elongated double-cone outer segments that enhanced long-wavelength sensitivity under blue light when compared to white (Wagner & Kröger, 2000). It can be speculated that an increase in LWS expression may serve to enhance visual contrast against a short-wavelength-dominated background (Lythgoe, 1979; Carleton & Kocher, 2001). Alternatively, the preferential modulation of long-wavelength opsins may reflect constraints in plasticity for the short-wavelength opsins, potentially shaped by the evolutionary and environmental history of the species.

Green chromides in this study originated from the turbid waters of the Sundarbans, where intense attenuation and scattering of short-wavelength light enrich the ambient spectrum in longer wavelengths (Mookherjee et al., 2015; Jones et al., 2021; Carleton et al., 2020; Bertinetti & Torres-Dowdall, 2025). The influence of natal photic environments on opsin plasticity is well documented in bluefin killifish (*Lucania goodei*), where turbid-water populations show minimal modulation of opsin (short-wavelength opsins) expressions, when compared to clear-water populations (Fuller & Claricoates, 2011). Synthesizing the responses observed in *A. pulcher* and *L. goodei*, we hypothesize that long term habitat specialisation in turbid, long-wavelength-dominated waters may constrain plasticity in short-wavelength opsins of green chromides, while maintaining regulatory flexibility in long-wavelength opsins. Testing whether this asymmetric plasticity explains the response of green chromides under blue light will require comparative studies across wild populations originating from distinct photic environments. Green chromides provide an ideal empirical system to test this hypothesis because wild populations occupy habitats spanning severe turbidity gradients, ranging from clear waters in Ratnagiri (0.3-8 NTU), Maharashtra, India (western India) and the Thamirabarani River (19-13 NTU), Tamil Nadu, India (southern India) to highly turbid systems such as Chilika Lake (96-260 NTU), Odisha, India (eastern India) (Khandagale et al., 2022; Kani & Murugesan, 2014; Jally et al., 2021). Such comparisons across these environmental gradients will determine whether the static short-wavelength opsin expression observed here is a population-specific artifact of turbid-water exposure or an invariant species-level trait reflecting the evolutionary lineage of green chromides.

Alongside the upregulation of LWS, RH2A expression decreased under blue light. Although comparison with the 90-day white-light control (W90) did not entirely separate out the spectral effects from temporal duration, but absolute values of RH2A transcript levels at B45 were lower relative to both W45 and W90, indicating a consistent light-induced down-regulation. The reciprocal expression trajectories of LWS and RH2A likely reflect altered coexpression ratios within double-cone photoreceptors. Coexpression of RH2 and LWS in double cones is well documented in cichlid, *Metriaclima zebra*, wherein relative transcript ratios shift dynamically in response to spectral manipulation (Dalton et al., 2014, 2015). Thus, green chromides may modulate double-cone spectral sensitivity by tuning the relative proportion of LWS and RH2A visual pigments within individual outer segments. Validating this cellular mechanism will require transcript localization via fluorescence in situ hybridization (FISH) or direct spectral profiling of individual double cones using microspectrophotometry (MSP) (Dalton et al., 2014, 2015).

Crucially, we showed that despite the apparent functional mismatch between ambient blue light and long-wavelength-biased opsin shifts, visually mediated behaviors remained entirely stable in our study species. Green chromides maintained strong conspecific preference at the immediate onset of blue-light exposure (B0) and displayed invariant latency in food detection throughout the experiment, showing no behavioral lag prior to or alongside the significant elevation in LWS transcript levels observed at Day 15 (B15). Furthermore, subsequent shifts in opsins (LWS and RH2A) and Cyp27c1 expression caused no measurable behavioral disruption, demonstrating that these social and foraging tasks are highly resilient to spectral alterations. This discrepancy in behavioural expressions and opsin expression patterns, have also been previously demonstrated in a range of different fish species, including both cichlids and non-cichlids (Fuller et al., 2010; Wright et al., 2020; Potkamp et al., 2026). Most of these studies have speculated that behavioural expressions, for example, mate choice or foraging, are highly complex in nature, which depend on multiple processing steps downstream of initial photon capture, thus buffering behavioral execution against peripheral retinal changes (Potkamp et al., 2026). However, we would like to highlight that participating in behavioral trials can itself alter retinal opsin profiles (Wilwert, 2022), and that too, in a time-dependent manner (Potkamp et al., 2026), for example, expressions during the trials versus after the trials. Evaluating whether behavioral testing influenced opsin expressions in the green chromides would require comparing trial-subjected fish against untested controls, a comparison we could not perform due to sample size constraints, leaving open the possibility of assessing test-induced effects on opsin expressions. Further the specific visual cues used by green chromides for conspecific recognition remain still unknown (e.g., whether they rely primarily on achromatic luminance, body morphology, motion patterns, or chromatic cues), we speculate behavioral performance may have been maintained through non-chromatic visual channels that are less sensitive to retinal opsin shifts.

Although plasticity in opsin expression is well documented across cichlids and broader vertebrate taxa (Hofmann et al., 2010; Nandamuri et al., 2017; Schott et al., 2022; Carleton et al., 2020), our findings in green chromides demonstrate that the interplay between the opsin gene expression and behavioral output is non-linear and complex. This disconnect underscores the intricate nature of the proximate mechanisms governing performances in visual tasks. Future studies should focus on unravelling the specific molecular pathways mediating social preferences in green chromides while explicitly defining the visual cues involved in conspecific recognition.

## Acknowledgement

We thank Dr. Rajesh Kushawaha, Associate Professor, Physics Research Laboratory, for his assistance with irradiance measurements. We are grateful to Hemadri Maity and Tapan Maity for providing the fish used in this study. CD acknowledges the University Grants Commission (UGC), Government of India, for fellowship support.

## Author Contributions

CD was responsible for conceptualization, data curation, formal analysis, investigation, methodology, validation, visualization, writing-original draft, and writing - review & editing AB was responsible for data curation, formal analysis and methodology for bioinformatics.

R. Ganguly was responsible for data curation for behaviour data. KBS contributed to conceptualization, formal analysis, supervision, writing - review & editing. R.Ghosal contributed to conceptualization, formal analysis, funding acquisition, methodology, project administration, resources, supervision, writing - review & editing.

## Data Availability statement

All data supporting this study, including irradiance measurements, gene expression, behavioural observations, and bioinformatics analyses, are available on Figshare [10.6084/m9.figshare.34883226]

## Declaration of generative AI and AI-assisted technologies in the manuscript preparation process

During the preparation of this work, the author(s) used ChatGPT, Google Gemini, and Grammarly for checking any grammatical errors. The author(s) reviewed and edited the output as needed and take full responsibility for the content of the published article.

