## Supplementary Material for "Social ties are maintained through sensory adaptations in green chromides"

### Table S1: Primer sequence and efficiency of each primer pair for each gene

| Gene | Forward primer | Reverse primer | Primer efficiency (%) | Slope |
|---|---|---|---|---|
| RH1 | ATGAATGGCACAGAGGGAC | CGTTGTGGTGAATCCTCCAA | 91.18 | -3.55 |
| SWS1 | CCACCTTTCTTTGGCTGGA | CAAAGTAAAGAGCTGCCAG | 96.34 | -3.41 |
| SWS2A | CGATCCCACCTCAACTACAT | TGAATGCACAGCAGGCTA | 100 | -3.3 |
| SWS2B | TCCCTCTGGACACCAACAA | CCAAGTTCACCAGGATGTAGT | 96.45 | -3.41 |
| LWS | CGTCACCCTCTAAACTGGAT | TGTCAGGGACCAAAGAGCA | 92.56 | -3.51 |
| RH2A | GCAAGAACTTCTACATTCCC | CACATGATGAGTCCAGCCA | 87.02 | -3.37 |
| EF1$\alpha$ | GTGAGCGCCTACATCAAGAA | CCAATCTTGTAGACGTCCTGC | 98.63 | -3.36 |
| UBE2Z | CAGCAGTGCATCCTGAGGATAAAG | TGATGAGCTTGACCCGTGG | 92.31 | -3.52 |
| GAPDH | GTGGGCAAAGTCATTCCTGA | GTGTAACCCAGCACTCCT | 89.31 | -3.61 |
| Cyp27c1 | CTGCTCACACACATGCTC | ATTGCCTGGGAGAACTGG | 84.78 | -3.75 |

### Table S2: Nucleotide sequences of each identified gene coding sequence (opsins and housekeeping) from green chromide

| **Gene** | **Sequence** | **Accession number** |
|---|---|---|
| RH1 | ATGAATGGCACAGAGGGACCCTTTTTTTATGTCCCTATGGTAAACACCACTGGTGTTGTTCGGAG<br>TCCTTATGAATACCCCCAGTACTACCTTGTCAACCCAGCAGCTTATGCTGCTCTGGGTGCCTATAT<br>GTTCTTTCTCATCCTTGCCGGCTTCCCCATCAACTTCCTTACTCTCTACGTCACTATCGAACACAA<br>GAAGCTGCGAACCCCACTAAACTACATCCTGCTGAACCTTGCAGTGGCTAACCTCTTCATGGTG<br>TTTGGAGGATTCACCACAACGATGTACACCTCTATGCATGGCTACTTTGTCCTAGGTCGCCTTGG<br>CTGCAATCTGGAAGGATTCTTTGCTACCCTCGGTGGCGAGATGTCGCTCTGGTCACTTGTTGTTC<br>TGGCTATTGAAAGGTGGGTGGTTGTCTGCAAGCCCATTAGCAACTTCCGTTTTGGGGAGAACCA<br>CGCAATTATGGGCCTGGCTTTCACCTGGACAATGGCCTTGTCTTGTGCTGGACCCCCTCTTGTTG<br>GCTGGTCACGTTACATCCCTGAGGGCATGCAGTGCTCATGTGGAGTTGACTACTACACACGTGC<br>AGAAGGTTTCAACAATGAGTCCTTCGTTATCTACATGTTCATTTGCCACTTCATCATTCCACTGAG<br>TGTTGTGTTTTTCTGCTATGGCCGCCTGCTCTGCGCTGTCAAGGAAGCTGCTGCTGCCCAGCAG<br>GAGTCTGAAACCACCCAGAGGGCTGAGAGGGAAGTCACCCGCATGGTTGTTATCATGGTCATTG<br>CTTTCCATATATGTTGGTTACCCTATGCAGGAGTGGCCTGGTGGATCTTCACACATCAAGGCTCTG<br>AATTTGGACCAATCTTGATGACCCTCCCAGCCTTCTTTGCCAAGAGTTCTTCCATCTACAACCCT<br>TTGATTTACATCTGCATGAACAAGCAGTTCCGCCACTGCATGATCACCACCTTGTGCTGTGGGAA<br>AAATCCCTTTGAAGAAGAGGAGGGAGCATCCACTACTGCCTCCAAGACCGAGGCTTCCTCTGCC<br>TCCTCCAGCTCTGTGTCTCCTGCATAA | PV849961.1 |
| LWS | ATGGCAGAAGAGTGGGGAAAACAGTCTTTTGCTGCTAGGCGGTACCATGAAGATACAACAAGG<br>GGATCGGCTTTCGTTTACACAAACAGCAATAACACCAGAGATCCCTTTGAGGGTCCCAATTACC<br>ATATTGCTCCTCGATGGGTTTACAATCTTGCAACACTATGGATGTTTGTTGTGGTCATCTTATCAGT<br>CTTCACTAATGGTCTCGTCATAGTGGCCACACTAAAGTTCAAGAAACTGCGTCACCCTCTAAACT<br>GGATCCTGGTCAATCTTGCAATTGCTGATCTGGGAGAAACAGTTTTTGCCAGCACAATTAGTGTA<br>TGCAACCAGTTTTTTGGCTACTTCATTCTGGGACACCCAATGTGTGTCTTTGAGGGCTACGTTGT<br>CTCAACTTGTGGTATTACTGCTCTTTGGTCCCTGACAATCATCTCCTGGGAAAGATGGATAGTTG<br>TGTGCAAACCTTTTGGAAATGTCAAGTTTGATGCCAAATGGGCCACAGGTGGAATTGTGTTCTC | PV849956.1 |

| | | |
|---|---|---|
| | CTGGGCCTGGGCAGCAGCATGGTGTTCTCCTCCCATCTTTGGATGGAGCAGGTACTGGCCTCAT<br>GGACTGAAAACCTCCTGCGGACCTGATGTATTCAGTGGAAGTGAGGACCCTGGAGTTCAGTCTT<br>ACATGATTGTCCTCATGATTACTTGTTGCATCATCCCCCTGGCTATCATCATCCTGTGCTACCTTGC<br>TGTGTGGTTGGCCATCCGTGCTGTTGCCATGCAGCAGAAGGAATCAGAGTCAACCCAGAAAGCT<br>GAGAAAGAAGTCTCCAGAATGGTGGTTGTCATGATCCTGGCTTATTGTGTGTGCTGGGGACCTT<br>ACACCTTCTTTGCCTGCTTTGCTGCGGCTAACCCTGGATATGCCTTCCATCCTCTGGCTGCTGCCA<br>TGCCTGCATACTTTGCCAAGAGCGCTACTATCTACAACCCAATCATCTATGTCTTCATGAACCGAC<br>AGTTCCGCACATGCATCATGCAGCTCTTTGGCAAACAAGTGGATGATGGCTCTGAAGTATCCAC<br>ATCAAAGACAGAGGTTTCCTCTGTGGCTCCTGCATAA | |
| RH2A | ATGACCTGGGAAGGAGGAATTGAGCCCAATGGTACTGAGGGCAAGAACTTCTACATTCCCATGT<br>CCAACAGGACTGGAATTGTTAGAAGTCCTTTTGATTATCCACAGTATTACCTGGCAGACCCGATA<br>TTCTTCAAGCTCCTGGCTGTCTACATGTTCTTCCTGATCTGCACGGGAACTCCCATCAACGGCTT<br>GACATTGTTTGTAACTGCTCAGAACAAGAAGCTCCGGCAACCTCTTAACTACATTCTGGTCAAC<br>CTGGCTGTGGCTGGACTCATCATGTGCCTCTTTGGATTCACCATCACCATTACATCAGCTGTTAAT<br>GGCTACTTCATTCTTGGACCCACATTCTGTGCTATTGAGGGGTTCATGGCCACACTTGGAGGTGA<br>GGTTGCTCTCTGGTCACTGGTTGTTCTGGCTGTTGAGAGATACATTGTTGTCTGCAAACCCATGG<br>GAAGCTTCAAGTTCAGCGGGGCCCACGCTGGTGCCGGAGTACTTTTCACCTGGATCATGGCTTT<br>AGCTTGTGCTGCACCACCACTTTTTGGCTGGTCCAGGTATATTCCAGAGGGAATGCAGTGCTCCT<br>GCGGTCCTGACTACTACACTCTGGCTCCAGGCTTCAACAATGAGTCATATGTCATCTACATGTTT<br>GTTGTCCACTTCTTTGTTCCTGTCTTCATCATTTTCTTCACTTATGGAAGCCTTGTGATGACAGTC<br>AAAGCTGCAGCTGCACAGCAGCAGGACTCAGCTTCTACCCAGAAAGCTGAGAAGGAAGTGAC<br>CCGTATGTGTGTCTTGATGGTGTTGGGCTTCCTGGTAGCTTGGACACCATATGCCACTTTTGCTG<br>GTTGGATTTTCCTGAACAAGGGAGCTGCCTTCACTGCCCTGACTGCAGCCCTCCCTGCTTTCTTT<br>GCAAAGAGCTCTGCCTTGTACAATCCAGTTATCTATGTGCTGATGAACAAACAGTTCCGTAACTG<br>CATGCTAACTACTATTGGAATGGGTGGCATGGTGGAGGATGAGACTTCAGTTTCAACAAGCAAG<br>ACAGAGGTGTCCTCTGTGTCTTAA | PV849960.1 |
| SWS1 | ATGAGTGTTTATAAAGAGGCTGAAGCTGTTGGCTGTGGTAGAAAGCTCAGGCTCATCATGGGGA<br>AATATTTCCATCTCTATGAGAACATCTCCAAAGTGAGTCCCTTTGAGGGGCCCCAGTACTACCTG<br>GCTCCTGTGTGGGTCTTCTACCTACAGGCTGCCTTTATGGGCTTTGTCCTGTTTTGCGGTGCACC<br>TCTAAACTTTACTGTTATCTTAGCAGCCATGAAGTACAAGAAGCTCCGAGTCCCGCTCAACTATA<br>TTCTTATCAACATCTCCATCTCTGGCTTCATGTTCTCCACCTTCTCTGTTAGTCAGGTGTTCATTTC<br>CTCCCTGAAAGGTTACTACATCTTTGGGCACACAATGTGTGCACTGGAGAGTTCTTTGGGCTCTG<br>CTGCAGGTCTGGTGACCGGCTGGTCTTTGGCAGTACTGTCCGCTGAGAGGTATCTGGTCATCTGT<br>AAACCATTTGGATCCTTTAAGTTCAGCTCAAGTCATGCTTTGGGAGCGGTGGCCTTCACCTGGTT<br>CATGGGAATCGGCTGTGCATGCCCACCTTTCTTTGGCTGGAGCAGATATATCCCTGAAGGTCTGG<br>GCTGCTCCTGTGGACCTGACTGGTACACTCACAGTGAGGAGTACAAGACCTCGAGCTACACTCA<br>CTTCCTAATGGTCACCTGCTTTATCATGCCTCTGACCATCATCATTTTCTCCTACTCCCAGCTGCTG<br>AGTGCGCTAAGAGCTGTTGCAGCCCAGCAGGCGGAGTCAGCATCCACTCAAAAGGCAGAGAA<br>GGAGGTATCGAAGATGATCATCGTGATGGTGGGATCCTTCATCACCTGCTACGGCCCGTATGCTC<br>TGGCAGCTCTTTACTTTGCCTACTCCCCAGATGAAAATAAAGATTACCGACTAGTCAGCATCCCA<br>GCGTTATTCTCTAAGAGCTCCTGCGTCTACAACCCACTGATCTATGCCTTCATGAACAAACAGTT<br>TAATGGCTGCATCATGGAGATGGTGTTTGGAAAGAAGATGGAGGATGCTTCTGAAGTTTCATCA<br>AAGACTGAAGTGTCTTCAGTATCCACAGCTTCCTAA | PV849957.1 |
| SWS2A | ATGAGGAGTAATCGTGGTATGGAGCTGCCAGAAGACTTCTGGATACCTGTCCCCTTGGACACAA<br>ACAACATCACAGCACTTAGCCCTTTCCTGGTTCCACAGACCCACTTGGGGAGTTCAGCCACCTA<br>CTATGCTTTGGCCGCATTCCTGTTCTTCATATTTATAGGTGGCACTTTCATCAATATCCTCACCATT<br>GCATGCACCATCCAGTACAAGAAACTTCGATCCCACCTCAACTACATTCTGGTGAACTTGGCATT<br>GTCAAACCTTCTTGTGGCCTGTGTGGGTTCTTTCATGGCCTGCATCTCCTTTGCAAGTAGATATTT<br>CATCTTTGGTTCACTTGGATGCAAGATTGAAGGTTTTATGGCAACACTTGGTGGTATGGTAAGCT<br>TGTGGTCTCTTTCCGTGATAGCTTTTGAAAGATGGCTTGTGATCTGCAAGCCACTTGGTAACTTT<br>GTTTTCAAGCCCGACCATGCTCTAGCCTGCTGTGCATTCACCTGGGTTTTTGCATTAATTGCCTCA<br>GCTCCTCCACTGTTTGGATGGAGCAGGTATATCCCAGAAGGCTTACAGTGCTCCTGTGGACCAG<br>ACTGGTACACCACAAACAACAAATATAACAATGAGTCCTATGTGATGTTCCTTTTCTGCTTCTGC<br>TTTGCTTTTCCCTTCGCTGTGATTGTCTTCTGCTACACCCAGCTGCTCATTACACTAAAAATGGCA<br>GCCAAGGCACAAGCTGAGTCTGCCTCCACCCAGAAGGCAGAGAAGGAAGTGACGAGGATGGT<br>GGTGGTCATGGTACTGGGCTTCCTGGTGTGCTGGTTGCCATACACCAGCTTTGCTCTTTGGGTTG<br>TGAACAATCGTGGCCAACAATTTGACCTGAGACTGGCAACTATACCCTCCTGCTTCTCGAAGTCT<br>TCCACAGTCTACAACCCTATCATCTATGTTCTCCTCAATAAACAGTTCCGTACATGTATGTTATCA<br>ATGCTGGGGATGGGAGGAGGTGAAGAAGAAACCTCAACAACACAATCAGTGACTGAAGTTTCC<br>AAAGTTGGCCCTGCTTAG | PV849959.1 |
| SWS2B | ATGAGGGGAAAGCAACCTGCAGAATTCCCAGATGACTTCTGGATCCCCATCCCTCTGGACACCA<br>ACAACATCACAGCTCTCAGTCCATTCCTGGTTCCCCAGGACCATTTAGGGAGCCCGTTCATCTTT<br>TATTCAATGTCAGCATTCATGTGCTTCTTGTGGGTGACTGGTACAGCTATCAACATCCTGACTATT<br>GTATGTACTATTCAATACAAGAAACTCCGGTCTCATCTGAACTACATCCTGGTGAACTTGGCTGT<br>GGCAAACCTCCTCGTCTCCTGCGTGGGCTCATTCACCTGCTTCATCTGTTTTGCCTTCAAATACAT<br>GTTTTTTGGTCCAGTGATGTGCAAGATTGAAGGATTTGCAGCAACTCTTGGTGGCATGGTTAGTC<br>TGTGGTCTCTTGCTGTGGTAGCATTTGAGAGATGGCTGGTTGTCTGCAAGCCACTTGGGAATTTT<br>GCCTTCAAGTCCAGCCATGCTATAGCATGCTGTGGATTGACTTGGGTCTTTGCTATGAGTGCTGC<br>TCTTCCTCCGCTGCTTGGATGGAGCAGGTATATCCCAGAAGGCTTGCAATGTTCCTGCGGCCCAG<br>ACTGGTACACAACAGACAACAAGTTTAACAATGAATCCTATGTGATTTTCCTCTTCTGCTTCTGC<br>TTTGCTGTCCCTTTCTGCGGCATCATCTTCTGCTACTCACAGCTACTTTTTATGCTAAAATCGGCA<br>GTGAAGGCCCAAGCTGAATCTGCCTCCACTCAGAAGGCGGAGAAGGAGGTGACCAGGATGGTG<br>ATCATCATGGTGCTGGGCTTCCTGGTGTGCTGGTTGCCATACACCACCTTTGCCCTTTGGGTTGT<br>GAACAATCGTGGGCAGCCATTCGACCTGAGACTGGCCACCATACCCTCCTGCCTCTCAAAGGCC<br>TCCACAGTCTACAACCCTGTCATCTACATCCTCCTCAATAAGCAGTTTCGTTCATGTATTAGGAAG<br>ATGCTGGGGATGAGTGGAGGCGACGACGAAGAATTATCAGCAACTCAATCAGTCACTGAAGTCT<br>CTAAAGTTGGACCTTCTTAG | PV849958.1 |

| | | |
|---|---|---|
| UBE2Z | ATGGCGGACAGCTTTGGAGAGGAGTCCAGCAGCGGTGCCCTTGGCTTGGACGTTACCGGACAG<br>GGCAGCGGAGCCTCGCTGCTGCCGATTTTGGCTAGCTCTCTGCCGGTGGGACACTCGGTAACCG<br>GCGGTGCTCACCCTAGCTCCAGCCCGGCCTCTCCTCCTCCTGCCGCGGCCGCGCAGAGTGGCCT<br>AACAGCTGTGGTGCCCCCTATGTCCGCTGTCACTGCTACTCCGGTCAGCTTTGGAAACACACTT<br>ACTCCCGGCTCCTCTCCAGCCGTCGTGGCCGTGTCTCCCACCGTGCATTCCTCCTCACCTCTGTC<br>CGGTGTTGGACTAGGGGTGGCCGGGGTGGCAGGAGCGGGCCTCCTGTCCCAAATCAACGCCAC<br>TTCCTGGGACCCAACTCTGAGTACGGACTGGGATAACGAGAAGGCTTCCCAGCAGTGCATCCTG<br>AGGATAAAGCGGGATATTATGTCCATCTACAAGGAACCTCCACCAGGGATGTTTGTCGTCCCTGA<br>CCCTCAAGACATGACCAAGATCCATGCTCTGATCACAGGACCATTCGACACACCATACGAGGGT<br>GGCTTCTTCCTCTTCCTGTTCCGCTGCCCCCCTGACTACCCTATTCACCCACCACGGGTCAAGCT<br>CATCACTACCGGACACAACACTGTCCGCTTCAACCCCAACTTCTACCGCAACGGCAAAGTATGC<br>CTCAGCATTCTGGGGACATGGACAGGCCCAGCATGGAGTCCAGCTCAGAGCATCTCATCTGTGC<br>TCATCTCCATCCAGTCTCTGATGACTGAGAACCCCTATCACAATGAACCTGGTTTTGAGCAGGAG<br>CGTCACCCAGGGGACAGTAAGAATTATAATGAGTGTATCCGCCATGAAACCATGAGGGTGGCAG<br>TGTGTGACATGCTGGAGGGCAAAGTTCCCTGTCCTGAAGCTTTGTCGAGTGTGATGGAAAAGTC<br>TTTCCTTGAATATTATGATTTCTACGAGGGAGTCTGCAAAGAGAGGCTGCATCTGCAGGGTCAGA<br>ACATGCAGGACCCATTTGGCGAGAAGCGTGGCCGCTTCGACTACCAGGGACTGCTGGCTCGCCT<br>CAGTGCAACCCACAGGCGAATACGGGAGAAAAACTTGGCAGAGGACGACCACAATGACAACG<br>ACTCAGACTCCGACACCAGCTCCTCAGGGACGGACCCTGACAGTCAGGGCAGCTCCCAGCCCT<br>GA | PV849963.1 |
| EF1α | ATGGGAAAGGAAAAGACCCACATTAACATCGTGGTCATTGGCCATGTCGACTCTGGCAAGTCCA<br>CCTCCACCGGCCACCTGATCTACAAGTGCGGAGGAATCGACAAGAGAACCATCGAGAAGTTCG<br>AGAAGGAAGCCGCCGAGATGGGAAAGGGCTCCTTCAAGTATGCCTGGGTGCTGGACAAACTGA<br>AGGCTGAGCGGGAGCGTGGTATCACCATTGACATCGCTCTGTGGAAGTTTGAGACCAGCAAGTA<br>CTATGTGACCATCATTGACGCCCCTGGACACAGGGACTTCATCAAGAACATGATCACTGGTACCT<br>CTCAGGCTGACTGTGCTGTGCTGATCGTCGCTGCTGGTGTTGGTGAGTTTGAGGCTGGTATCTCC<br>AAGAACGGCCAGACCCGTGAGCACGCCCTGCTGGCCTTCACCCTGGGTGTGAAGCAGCTCATT<br>GTTGGAGTGAACAAGATGGACTCCACTGAGCCCCCCTACAGCGAGGCCCGTTTCAACGAAATC<br>ACCAAGGAAGTGAGCGCCTACATCAAGAAGATCGGCTACAACCCCAATGCTGTTGCTTTCGTCC<br>CCATCTCTGGATGGCACGGAGACAACATGCTGGAGGCCAGTGACAAGATGCCCTGGTTTAAAG<br>GGTGGAAAATTGAGCGTAAGGAGGGCAATGCCTCTGGAACCACACTGCTGGAGGCTCTGGATG<br>CCATCCTGCCCCCCGCCCGCCCCACCGACAAGCCCCTGCGTCTGCCCCTGCAGGACGTCTACAA<br>GATTGGGGGTATTGGAACCGTACCCGTCGGTCGTGTTGAGACTGGCATCCTGAAGCCTGGCACA<br>GTTGTCACCTTTGCTCCTGTTAACCTGACCACTGAGGTGAAGTCTGTGGAGATGCACCACGAGT<br>CTCTGCCCGAGGCTGTCCCTGGTGACAATGTTGGCTTCAACGTCAAGAACGTGTCCGTCAAGGA<br>AATCCGTCGTGGATACGTCGCTGGAGACAGCAAGAACGACCCACCCAAGGGTGCTGAGAGCTT<br>CAACGCTCAGGTCATCATCCTGAACCACCCTGGCCAGATTGGTCCAGGCTACGCCCCAGTGCTG<br>GATTGCCACACCGCCCACATTGCCTGCAAGTTCAGCGAGCTCATTGAGAAGATTGATCGTCGTT<br>CTGGCAAGAAGCTTGAGGACAGTCCCAAGTTTGTCAAATCTGGAGATGCCGCCATCGTCAAACT<br>GCTCCCACAGAAGCCCATGGTTGTGGAGCCTTTCTCCAGCTATCCTCCCCTTGGTCGTTTTGCTG<br>TGCGTGACATGAGGCAGACAGTGGCTGTTGGTGTCATCAAGTCTGTTGTCCCCAAGGAAGTTAG<br>CGGAAAGACAACCAAGGCTGCTGAGAAGGCCCAGAAGGGCAAGAAATGA | PV849964.1 |
| GAPDH | ATGAAGCCTGCTGAAATCCCCTGGGGCAGCTCTGGAGCCAAATATGTTGTCGAGTCCACTGGAG<br>TGTTCCTCAGTATCGAGAAGGCTTCTGCTCACCTGCAGGGTGGTGCGCAGCGTGTGGTTGTGTC<br>AGCCCCCTCACCTGATGCTCCAATGTTCGTCATGGGAGTTAATGAGGACAAATATGACCCTTCCT<br>CCATGACCATTGTCAGTAATGCCTCCTGCACCACCAACTGCCTGGCCCCCCTGGCCAAAGTCATC<br>CATGATAACTTTGGCATCGAGGAAGCCCTCATGACAACAGTCCACGCATACACAGCCACACAGA<br>AGACAGTGGATGGTCCCAGTGCCAAGGCCTGGCGTGATGGCCGTGGTGCCCATGCAAACATCAT<br>TCCAGCTTCTACCGGTGCTGCCAAGGCAGTGGGCAAAGTCATTCCTGAACTCAATGGTAAGCTG<br>ACAGGTATGGCATTCAGGGTGCCAGTGGCTGACGTGTCAGTGGTCGACCTGACATGCCGTCTTT<br>CCAAGCCTGCATCATACGCTGATATTAAGGAAGCTGTCAAGAAGGCCGCACAGGGACCCTTGAA<br>AGGAGTGCTGGGTTACACTGAGGACCAGGTGGTGTCCGCTGACTTTATCGGTGATACCCATTCC<br>TCGATTTTTGATGCTGGTGCTGGCATCTCTCTCAACGACAACTTTGTCAAGCTCATTTCCTGGTAT<br>GATAACGAGTATGGCTACAGCAACCGTGTCGCTGACTTGCTGCAGTACATGAACTCCAAGGAGT<br>AG | PV849962.1 |
| Cyp27c1 | ATGACGACCAAGCGTTTTGGGAACGCGTGCTGGAGAAGTTTTCAAGCCGGTCGGCTCAACGCG<br>CAGCTGTTCTACGTCGTGCGGACCCTGCACAAGTCTGCAGCCAGCGAGGCTTTGCGAATCTCCG<br>CGACCGGAGAGGAGGCCATACCCCAGGGGCTGATCAAGCCGGCAGATGTGGTCGAGAACACGA<br>CTATCAAGACCCTGAAGGAGATGCCCGGACCGAGCACCCTGTCCAACCTGATCGAGTTCTTCTG<br>GAAGGACGGGTTTAGCAGAATCCATGAAATTCAGCTGGAGCACCGGAAGAAGTATGGAAAAAT<br>TTTCAAGTCTCGTTTTGGGCCCCAGCTGGTGGTGTCCATTGCAGACCGTGACATGGTGGCCCAG<br>GTCCTGAGGGCTGAGGGCGTGGCCCCTCAGAGAGCTAACATGGAATCCTGGCAGGAATACAGA<br>GATATGAGAGGCCGGTCCACTGGCCTGATCTCAGCTGAGGGAGATGATTGGCTGAAGATGCGGA<br>GCGTGCTCAGACAGCTCATCATGCGGCCCCGTGATGTAGCAGTCTTCTCTGATGATGTAAACAAC<br>GTGGTAGATGACCTGATTGAAAGGGTCCGTAGTCTGCGTTCCCAGCAGGCTGACGGCGCAACCG<br>TCCTCAATGTGAATGACCTCTTCTTCAAATATGCCATGGAAGGTGTGGCTGCTCTTCTGTATGAAT<br>GCAGACTAGGATGTTTGGAAAATAATGTTCCCCAGAAAACCCAGGACTACATTGCTGCACTGAA<br>CCTCATGTTCAGCTCCTTTAAGACAACCATGTATGCCGGAGCGATCCCCAAGTGGCTACGACCTG<br>TCATCCCCAAACCATGGGAAGAGTTCTGTCACTCCTGGGATGGCCTGTTTAAATTCAGCATTATT<br>CATGTTGAGAAGAGATTCTCAGAGATCAACGCCAAGCTGCAGCGGGGAGAGGAGGTGAAGGG<br>GGGGCTGCTCACACACATGCTCATTACTAAGGAGATGAGCATGGAGGAGATCTACGCAAACGTA<br>ACAGAGATGCTGCTGGCTGGGGTTGACACGACATCATTTACACTGTCTTGGGCTACCTACCTGTT<br>AGCCCGTCACCCTCATATACAGCAGCAAATCTTTACAGAAGTTAATAGGACTCTGGGACCTGGA<br>GCAGTCCCCACAGCTGAGGATGTCCCTCGTCTGCCTCTCATCAGAGGGCTGGTCAAAGAGACTC<br>TCAGGCTTTTTCCAGTTCTCCCAGGCAATGGACGGATTACTCAGGATGACTTAGTGGTCGGTGG<br>ATACTTCCTTCCAAAAGGGACTCAGTTGGCCCTGTGTCACTACTCCACATCTTTGGAAGAGGAG<br>AACTTTTCTGATGCTCTGGACTTCCAGCCAGATCGCTGGATAAGGAAAGATTCCTCAGATCGTGT<br>CGACAACTTTGGCTCCATTCCTTTTGGCTATGGCATTAGGAGCTGCATTGGCAGGAGAATAGCAG<br>AGCTGGAGATGCACTTAGCTCTCACAAGGCTCATTCAGAAATTCCAGATTCGCTTGTCTCCTGTC<br>ACTGATGACGTCAAGGCCAAAACCCACGGCCTGCTCTGCCCTGCTGCCCCAATCCACCTGCAGT<br>TCGTTGACAGGGAACTAGATGCATGA | QB173709 |

**Table S3: Mean values of transcripts per million (TPM) for all the opsin genes and Cyp27c1 gene derived from the RNA-sequencing data**

| Gene | Mean TPM (transcripts per million) |
|---|---|
| RH1 | 23181 |
| LWS | 5256.41 |
| RH2A | 2448.13 |
| SWS1 | 180.81 |
| SWS2A | 1063 |
| SWS2B | 7.11 |
| Cyp27c1 | 7.15 |

**Table S4: Results of quasibinomial GLM model (expression~day*light) for dependent normalization of LWS, RH2A, SWS2A, SWS2B, and SWS1 expressions.**

| Predictor | Estimate | SE | t value | p-value |
|---|---|---|---|---|
| LWS | | | | |
| (Intercept) | 0.57778 | 0.07874 | 7.338 | 1.41e-07 |
| LightWhite | 0.11406 | 0.11234 | 1.015 | 0.3201 |
| DayDay15 | 0.33883 | 0.11487 | 2.950 | 0.0070 |
| DayDay30 | 0.27525 | 0.11408 | 2.413 | 0.0238 |
| DayDay45 | 0.36186 | 0.11518 | 3.142 | 0.0044 |
| LightWhite:DayDay15 | -0.42629 | 0.16082 | -2.651 | 0.0140 |
| LightWhite:DayDay30 | -0.31770 | 0.16052 | -1.979 | 0.0594 |
| LightWhite:DayDay45 | -0.47592 | 0.16090 | -2.958 | 0.0069 |

| | | | | |
|---|---|---|---|---|
| Dispersion | 0.00571 | | | |
| RH2A | | | | |
| (Intercept) | -1.61722 | 0.07792 | -20.754 | <2e-16 |
| LightWhite | -0.20212 | 0.11428 | -1.769 | 0.0897 |
| DayDay15 | -0.19029 | 0.11402 | -1.669 | 0.1081 |
| DayDay30 | -0.14627 | 0.11308 | -1.294 | 0.2081 |
| DayDay45 | -0.31569 | 0.11691 | -2.700 | 0.0125 |
| LightWhite:DayDay15 | 0.30547 | 0.16258 | 1.879 | 0.0725 |
| LightWhite:DayDay30 | 0.32139 | 0.16112 | 1.995 | 0.0576 |
| LightWhite:DayDay45 | 0.51781 | 0.16349 | 3.167 | 0.0042 |
| Dispersion | 0.0033 | | | |
| SWS1 | | | | |
| (Intercept) | -1.8880 | 0.1447 | -13.049 | 2.16e-12 |
| LightWhite | -0.1725 | 0.2116 | -0.815 | 0.4230 |
| DayDay15 | -0.3636 | 0.2207 | -1.648 | 0.1124 |
| DayDay30 | -0.4337 | 0.2243 | -1.933 | 0.0651 |
| DayDay45 | -0.3593 | 0.2204 | -1.630 | 0.1161 |
| LightWhite:DayDay15 | 0.3285 | 0.3115 | 1.054 | 0.3022 |
| LightWhite:DayDay30 | 0.5084 | 0.3110 | 1.635 | 0.1151 |
| LightWhite:DayDay45 | 0.5319 | 0.3056 | 1.740 | 0.0946 |
| Dispersion | 0.00096 | | | |
| SWS2A | | | | |

| | | | | |
|---|---|---|---|---|
| (Intercept) | -3.00097 | 0.11138 | -26.944 | <2e-16 |
| LightWhite | 0.40831 | 0.14509 | 2.814 | 0.0096 |
| DayDay15 | -0.24225 | 0.16700 | -1.451 | 0.1598 |
| DayDay30 | 0.14521 | 0.15262 | 0.951 | 0.3509 |
| DayDay45 | -0.00879 | 0.15782 | -0.056 | 0.9560 |
| LightWhite:DayDay15 | 0.38607 | 0.21020 | 1.837 | 0.0787 |
| LightWhite:DayDay30 | -0.47567 | 0.20857 | -2.281 | 0.0317 |
| LightWhite:DayDay45 | -0.39952 | 0.21438 | -1.864 | 0.0747 |
| Dispersion | 0.0022 | | | |
| SWS2B | | | | |
| (Intercept) | -4.1833 | 0.1222 | -34.232 | <2e-16 |
| LightWhite | -0.2683 | 0.1853 | -1.448 | 0.1605 |
| DayDay15 | -0.2343 | 0.1836 | -1.276 | 0.2140 |
| DayDay30 | -0.5486 | 0.2011 | -2.728 | 0.0117 |
| DayDay45 | -0.2302 | 0.1834 | -1.255 | 0.2214 |
| LightWhite:DayDay15 | 0.1288 | 0.2731 | 0.472 | 0.6414 |
| LightWhite:DayDay30 | 0.3566 | 0.2885 | 1.236 | 0.2285 |
| LightWhite:DayDay45 | 0.4985 | 0.2607 | 1.912 | 0.0678 |
| Dispersion | 0.0009 | | | |

**Table S5: Results of quasibinomial model (expression~treatment) on dependent normalisation of LWS and RH2A expressions in order to test the effect of time.**

| Predictor | Estimate | SE | t value | p-value |
|---|---|---|---|---|

| LWS | | | | |
|---|---|---|---|---|
| (Intercept) | 0.93964 | 0.05463 | 17.2 | 3.42e-08 |
| TreatmentW45 | -0.36186 | 0.07486 | -4.834 | 0.000929 |
| TreatmentW90 | -0.19314 | 0.07582 | -2.547 | 0.031325 |
| Dispersion | 0.00241 | | | |
| RH2A | | | | |
| (Intercept) | -1.93291 | 0.08406 | -22.994 | 2.65e-09 |
| TreatmentW45 | 0.31569 | 0.11276 | 2.800 | 0.0207 |
| TreatmentW90 | 0.21044 | 0.11461 | 1.836 | 0.0995 |
| Dispersion | 0.0031 | | | |

**Table S6: Results of quasibinomial GLM model (Preferences~Day*Light) for social preferences during green chromides versus tilapia trials**

| Predictor | Estimate | SE | t value | p-value |
|---|---|---|---|---|
| (Intercept) | 1.901 | 0.267 | 7.109 | <0.001 |
| LightWhite | 0.000 | 0.378 | 0.000 | 1.000 |
| DayDay15 | -0.316 | 0.359 | -0.879 | 0.382 |
| DayDay30 | -0.204 | 0.365 | -0.559 | 0.578 |

| | | | | |
|---|---|---|---|---|
| DayDay45 | 0.227 | 0.396 | 0.573 | 0.568 |
| LightWhite:DayDay15 | 0.000 | 0.508 | 0.000 | 1.000 |
| LightWhite:DayDay30 | 0.000 | 0.516 | 0.000 | 1.000 |
| LightWhite:DayDay45 | -0.355 | 0.542 | -0.655 | 0.515 |
| Dispersion | 0.081 | | | |

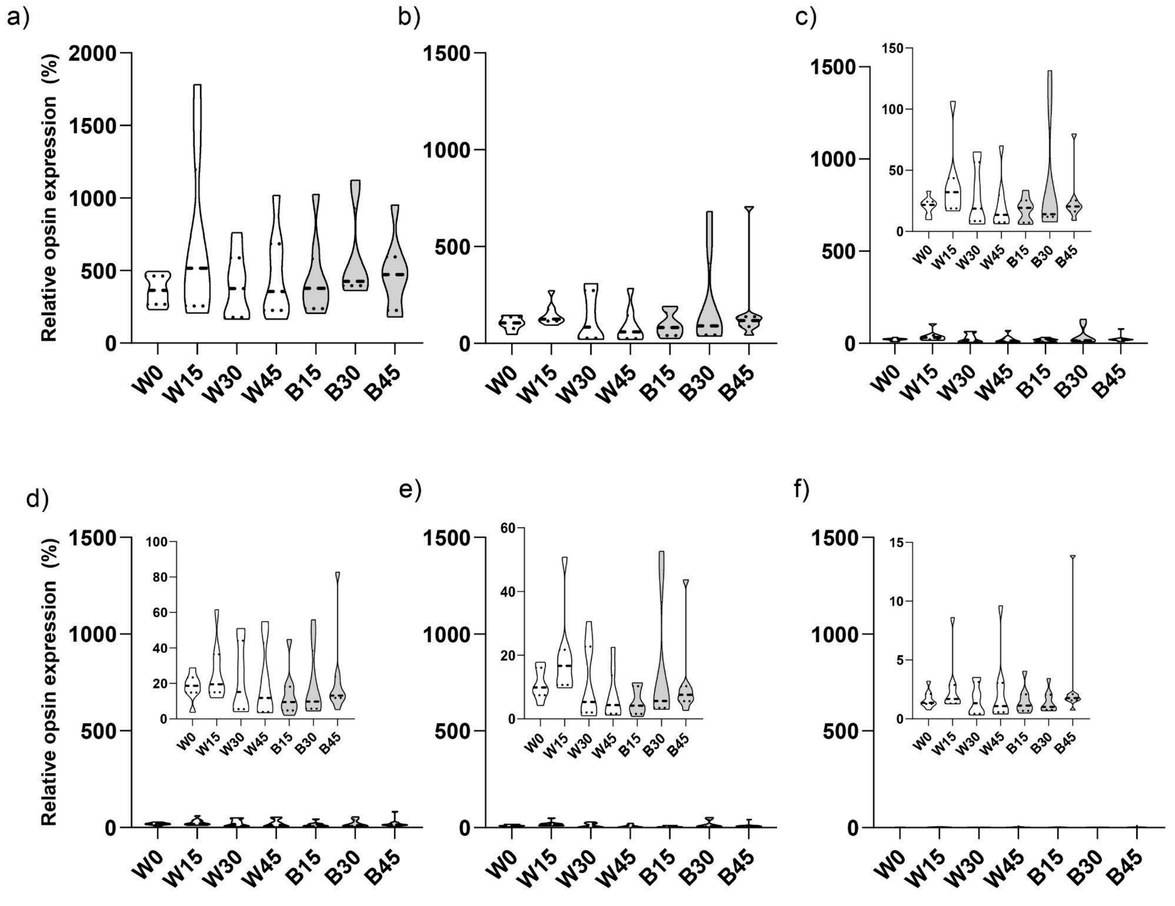

**Figure S1.** ***Independent normalisation of opsin expression for the entire experimental timeline including both white and blue lights***

*Violin plots illustrate relative expression (Ti/THk) of opsin genes, expressed as a percentage relative to EF1α. Panels show expression of (a) RH1, (b) LWS, (c) RH2A, (d) SWS1, (e) SWS2A, and (f) SWS2B genes. White and grey violins indicate white- and blue-light treatments, respectively. Each violin depicts the distribution of normalized expression values, with each individual contributing two data values (one from each of two independent plate runs). The dashed black line within each violin represents the median, while the dotted black lines represent the upper and lower quartiles. Insets within each panel present scaled representations of the same data to enhance data visualization.*

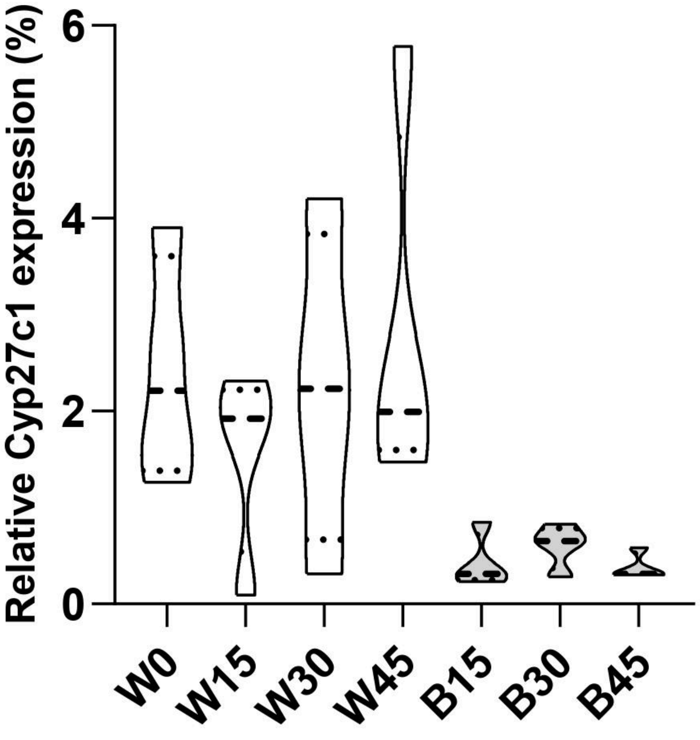

**Figure S2.** ***Independent normalisation of Cyp27c1 expression for the entire experimental timeline including both white and blue lights***

*Violin plots illustrate relative expression for Cyp27c1 gene. White and grey violins indicate white- and blue-light treatments, respectively. Each violin depicts the distribution of normalized expression values, with each individual contributing one data value (from a single plate run). The dashed black line within each violin represents the median, while the dotted black lines represent the upper and lower quartiles.*

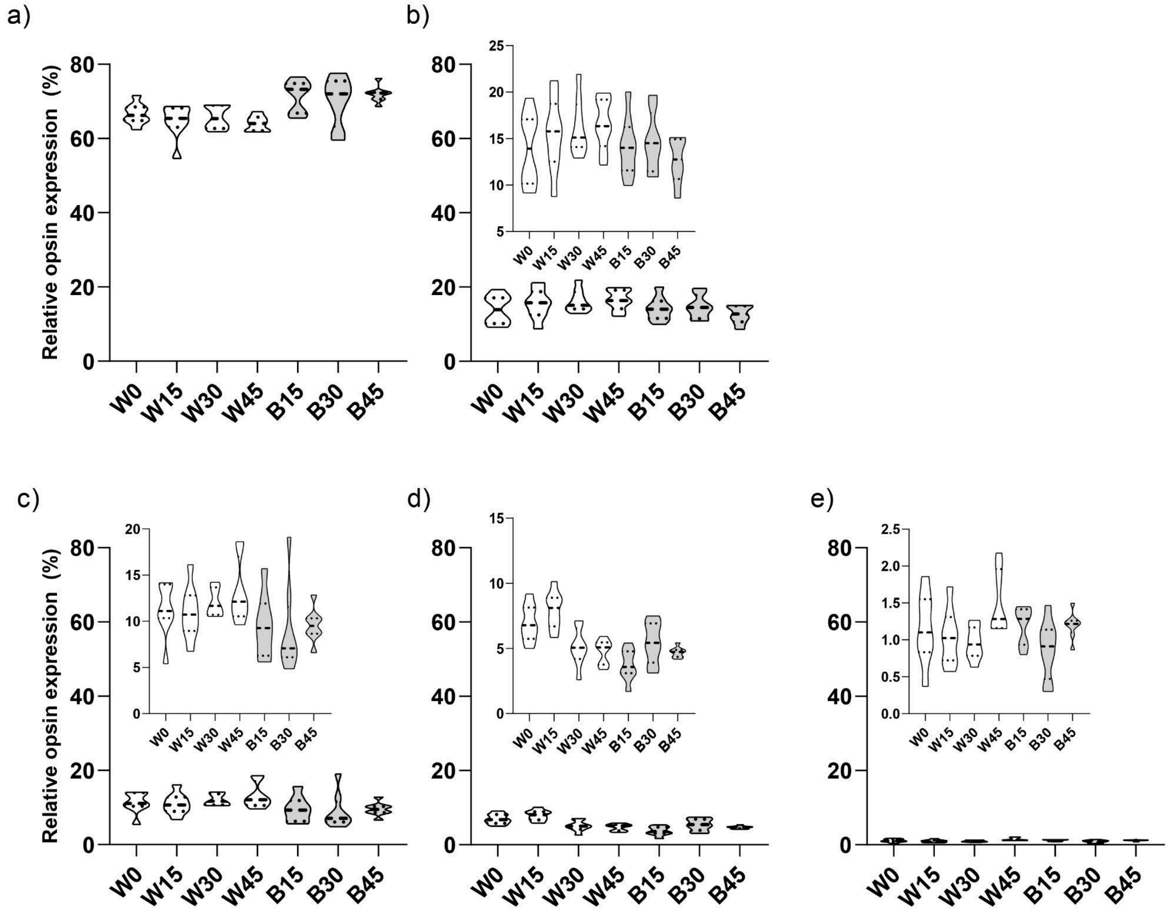

**Figure S3. *Dependent normalisation of opsin expression for the entire experimental timeline including both white and blue lights***

*Violin plots illustrate relative expression for each opsin gene expressed as a percentage. Panels show expression of (a) LWS, (b) RH2A, (c) SWS1, (d) SWS2A, and (e) SWS2B genes.*

*White and grey violins indicate white- and blue-light treatments, respectively. Each violin depicts the distribution of normalized expression values, with each individual contributing two data values (one from each of two independent plate runs). The dashed black line within each violin represents the median, while the dotted black lines represent the upper and lower quartiles. Insets within each panel present scaled representations of the same data to enhance data visualization.*

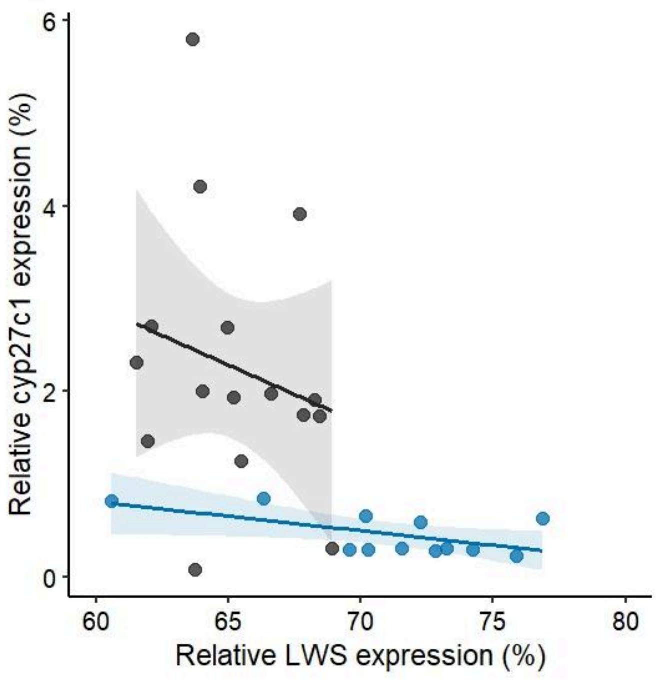

***Figure S4. Relationship between LWS and Cyp27c1 expressions.*** Black and blue symbols represent individual fish under white and blue lights, respectively. Shaded areas indicate ±95% confidence intervals.

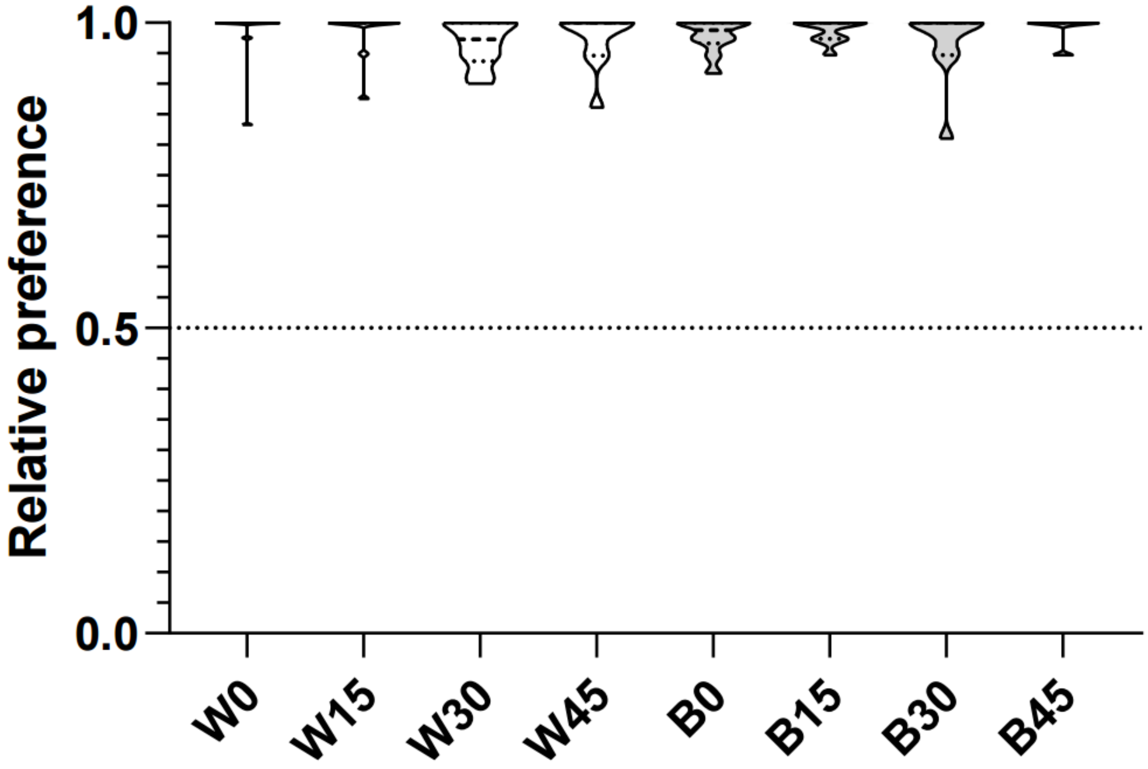

**Figure S5.** ***Social preferences measured during 0 versus green chromide trials***

*Violin plots illustrate social preferences of green chromides during 0 versus green chromide trials. White and grey violins indicate white- and blue-light treatments, respectively. The dashed black line within each violin represents the median, while the dotted black lines represent the upper and lower quartiles. A dashed line at 0.5 relative preference indicates random preference.*

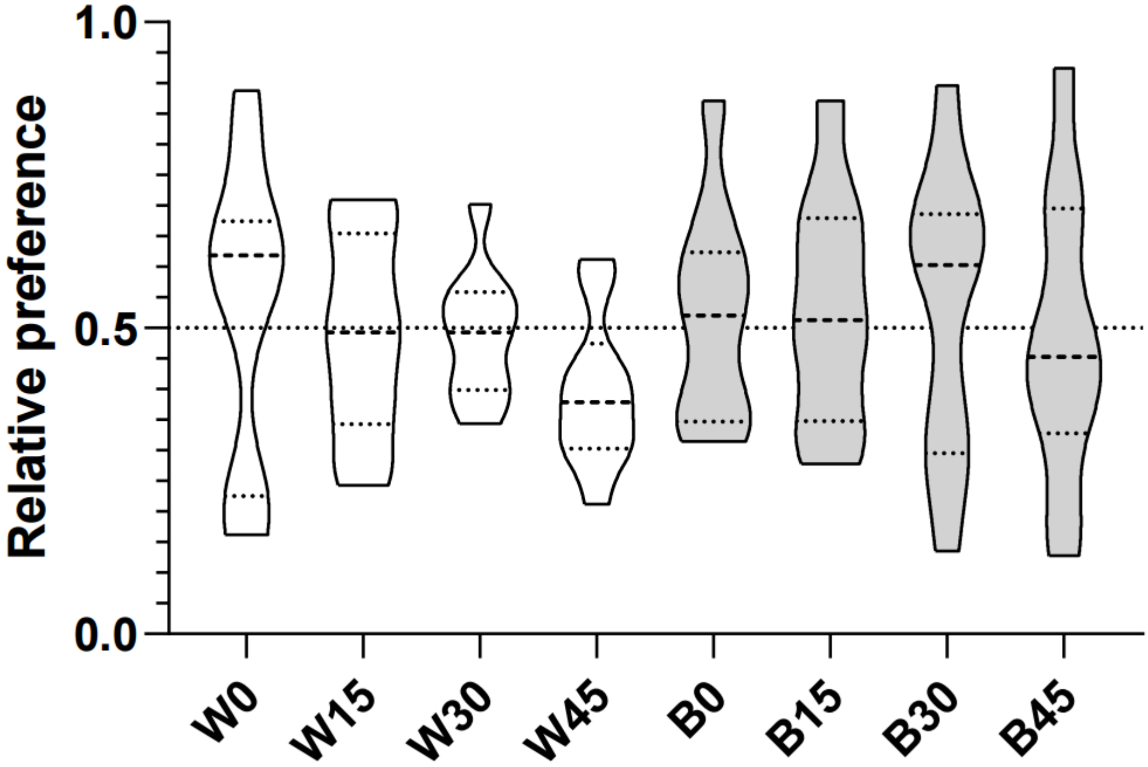

**Figure S6.** ***Social preferences of green chromides during 0 versus 0 trials***

*Violin plots illustrate social preferences of green chromides during 0 versus 0 trials. White and grey violins indicate white- and blue-light treatments, respectively. The dashed black line within each violin represents the median, while the dotted black lines represent the upper and lower quartiles. A dashed line at 0.5 relative preference indicates random preference.*

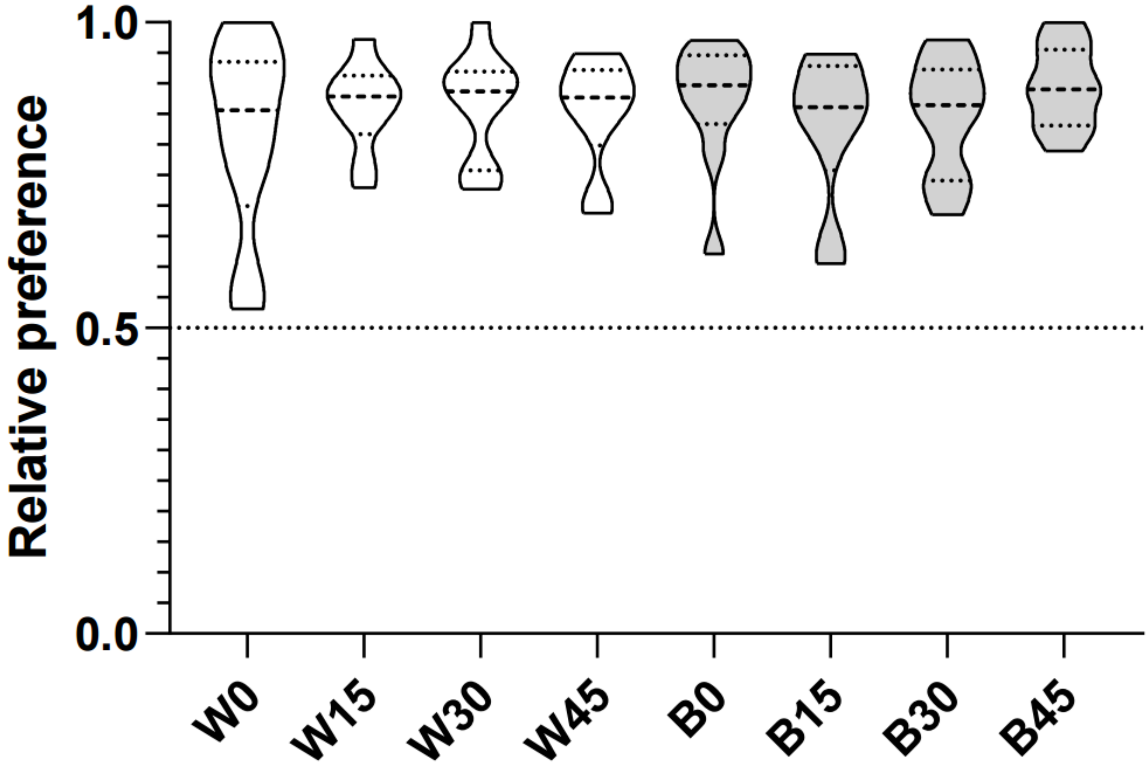

**Figure S7.** ***Social preferences of green chromides during green chromide versus tilapia trials***

*Violin plots illustrate social preferences of green chromides during green chromide versus tilapia trials. White and grey violins indicate white- and blue-light treatments, respectively. The dashed black line within each violin represents the median, while the dotted black lines represent the upper and lower quartiles. A dashed line at 0.5 relative preference indicates random preference.*
